# Disruption of membrane cholesterol organization impairs the concerted activity of PIEZO1 channel clusters

**DOI:** 10.1101/604488

**Authors:** P. Ridone, E. Pandzic, M. Vassalli, C. D. Cox, A. Macmillan, P.A. Gottlieb, B. Martinac

**Affiliations:** The Victor Chang Cardiac Research Institute, Lowy Packer Building, Darlinghurst, NSW 2010, Australia; Biomedical Imaging Facility (BMIF), Mark Wainwright Analytical Centre, Lowy Cancer Research Centre, The University of New South Wales, NSW, 2052, Australia; Institute of Biophysics, National Research Council, Genova, Italy; Physiology and Biophysics, State University of New York at Buffalo, Buffalo, NY, 14214, USA; St Vincent’s Clinical School, University of New South Wales, Darlinghurst, NSW 2010, Australia

**Keywords:** Mechanosensitive channel, Cholesterol, STORM, Patch-Clamp, Fluorescence Microscopy

## Abstract

The human mechanosensitive ion channel PIEZO1 is gated by membrane tension and regulates essential biological processes such as vascular development and erythrocyte volume homeostasis. Currently, little is known about PIEZO1 plasma membrane localization and organization. Using a PIEZO1-GFP fusion protein, we investigated whether cholesterol enrichment or depletion by methyl-β-Cyclodextrin (MBCD) and disruption of membrane cholesterol organization by dynasore affects PIEZO1-GFP’s response to mechanical force. Electrophysiological recordings in the cell-attached configuration revealed that MBCD caused a rightward shift in the PIEZO1-GFP pressure-response curve, increased channel latency in response to mechanical stimuli and markedly slowed channel inactivation. The same effects were seen in native PIEZO1 in N2A cells. STORM super-resolution imaging revealed that, at the nano-scale, PIEZO1-GFP channels in the membrane associate as clusters sensitive to membrane manipulation. Both cluster distribution and diffusion rates were affected by treatment with MBCD (5 mM). Supplementation of poly-unsaturated fatty acids appeared to sensitize the PIEZO1-GFP response to applied pressure. Together, our results indicate that PIEZO1 function is directly dependent on the membrane mechanical properties and lateral organization of membrane cholesterol domains which coordinates the concerted activity of PIEZO1 channels.

**SUMMARY:** The essential mammalian mechanosensitive channel PIEZO1 organizes in the plasma membrane into nanometric clusters which depend on the integrity of cholesterol domains to rapidly detect applied force and especially inactivate syncronously, the most commonly altered feature of PIEZO1 in pathology.

## INTRODUCTION

Mechanosensitive ion channels are membrane proteins that sense mechanical stimuli allowing cells to respond and adapt to physical forces. An essential family of eukaryotic mechanosensitive channels are PIEZO channels [1] comprised of two members, PIEZO1 and PIEZO2. These channels are associated with a number physiological functions such as the development of vascular architecture [2, 3]. Human PIEZO1 also plays a key role in erythrocyte volume regulation [4] where gain of function variants (GOF) cause Hereditary Dehydrated Stomatocytosis (Xerocytosis) ([5-8]). These mutations invariably slow channel inactivation and this phenotype confers native resistance to malarial invasion in African populations [9].

Three recently published cryo-EM structures of purified mouse PIEZO1 provide detailed information of its trimeric, curved structure [10-13]. Three homomeric subunits form a central non-selective cation permeable pore with a large-scale propeller-like assembly [10, 11]. Large beams that extend from the central pore axis to the extremity of the protein have been proposed to underlie a lever-like mechanism to transfer force from the periphery to the channel gate [11, 12]. Activation of PIEZO1 is associated with a force-from-lipids mechanism [14-18], where membrane forces are sufficient to drive a, yet unknown, conformational change in PIEZO1 and trigger gating. This suggests that both global effects on membrane physical properties and specific lipid interactions are likely to be essential for PIEZO1 function [19-21].

Cholesterol is a ubiquitous component of cellular membranes and a major modulator of membrane mechanical properties. Application of the cholesterol-depleting agent methyl-beta-cyclodextrin (MBCD) reduced whole-cell indentation induced currents in PIEZO1-expressing cells [22]. This response was linked to the cholesterol binding protein STOML3 previously shown to sensitize PIEZO1 channels [23]. Furthermore a crosslinking-based proteomics study reported a non-random degree of association between a cholesterol analog and PIEZO1 [24]. Thus, there is mounting evidence indicating physiologically-relevant interplay between cholesterol and PIEZO1-mediated mechanotransduction. In humans, the difference in the tissue expression pattern of STOML3 and the more ubiquitous PIEZO1 suggests that STOML3-mediated cholesterol sensitization of PIEZO1 may be tissue specific [25].

In this study we characterized the effects of cholesterol on PIEZO1 activity in cells with undetectable levels of STOML3 [25, 26]. Using super resolution (STORM) imaging we showed that PIEZO1 channels in the plasma membrane associate as clusters. Cholesterol removal or disruption affected the dynamics of these PIEZO1 clusters but had only a minor effect on their size. Removal of cholesterol or disruption of cholesterol rich domains reduced channel sensitivity, slowed down activation and abolished inactivation in cell-attached patches. We imaged both cholesterol-rich membrane domains and PIEZO1 in HEK293T cells stably expressing the channel protein, using both standard and super-resolution fluorescence microscopy to quantify the dynamic behavior of PIEZO1 and its relationship with cholesterol-rich membrane domains. Furthermore, we suggest a model of the PIEZO1 and cholesterol interaction and propose that this association is essential for the spatio-temporal activity of PIEZO1.

## METHODS

### Patch Clamp Electrophysiology

All recordings of human PIEZO1 were obtained from HEK293T cells stably expressing PIEZO1-1591-GFP [16], a previously characterized fusion protein [18]. N2A cells were a kind gift from Dr. K.Poole (University of New South Wales). PIEZO1 KO HEK293T cells were a kind gift from Prof A Patapoutian (Scripps Institute). Cells were plated onto 12-mm round glass coverslips and grown in Dulbecco’s Modified Eagle Medium (DMEM) supplemented with 10% Fetal Bovine Serum (FBS). Cells were treated with 5 mM Methyl-β-CycloDextrin (MBCD; C4555, Sigma), 100 μg/ml Water-soluble Cholesterol/MBCD complex (C4951, Sigma; product contains 44 mg Cholesterol per gram of compound) or 80 μM Dynasore (D7693, Sigma) in serum-free DMEM for 30 mins at 37°C. Untreated, control samples were also incubated in serum-free DMEM at 37°C for 30 min prior to patch-clamping. Docosahexanoic Acid (DHA) treatment was performed by culturing cells for 24 hr or 48 hr in DMEM supplemented with 10% FBS and 25 μM DHA (D2534, Sigma). All recordings were performed in identical pipette/bath solutions having the following composition: 140 mM NaCl, 3 mM KCl, 1mM MgCl_2_, 1mM CaCl_2_, 10mM Glucose, 10mM HEPES, pH 7.4. Borosilicate glass pipettes (Drummond Scientific, Broomall, PA) were pulled using a Narishige puller (PP-83; Narishige) to achieve the bubble number of 6.0 (3 MΩ pipette resistance). The PIEZO1 currents were recorded at room temperature using an AxoPatch 200B amplifier (Axon Instruments) in the cell-attached patch configuration, at a 20 kHz sampling rate with 1 kHz filtration. Negative pressure was applied to the patch membrane using a computer-controlled high-speed pressure clamp-1 apparatus (HSPC-1; ALA Scientific Instruments).

### Cell culture and sample preparation for imaging

PIEZO1-GFP stable HEK293T cells were seeded in DMEM (Gibco) supplemented by 10% FBS inside 35 mm diameter Fluorodishes (FD35-100, World Precision Instruments) and grown to 70-85 % confluency for 2-3 days.

For co-localization studies, cells were incubated in 1:1000 dilution of Cholera Toxin Subunit B (CtxB)-Alexa555 (Invitrogen, 5mg/ml stock) and rinsed 3 times with PBS. After labelling, treatments were performed as described above and cells were imaged live immediately after on a Zeiss Elyra microscope.

For STORM experiments cells were treated with drugs, 5 mM MBCD, 50 µg/ml water-soluble Cholesterol-loaded MBCD or 80 μM Dynasore diluted in serum-free DMEM, or incubated in serum-free DMEM for control samples for 30 min at 37 °C. Post treatment cells were washed with warm PBS and fixed with 4 % PFA for 15 min at room temperature followed with 3X rinsing with PBS. Post-fixation cells were permeabilized with Triton-X100 (0.01 % v/v) at room temperature for 15 min. After permeabilization cells were blocked with 5 % BSA for 1 h at room temperature, followed by incubating the dishes with labelling reagents at room temperature for 1 hr, PIEZO1-GFP was labelled with anti-GFP-Alexa-647 with 1:1000 PBS dilution from stock (Bioss Antibodies, 1 μg/μL). Samples were stored at 4 °C and imaged within 2 days from preparation on a Zeiss Elyra microscope.

### STORM imaging of Piezo-GFP-antiGFP-Alexa-647 and cluster point data analysis

STORM images were acquired on a total internal reflection fluorescence (TIRF) microscope (ELYRA, Zeiss) using a 100 × Alpha-Plan APO oil immersion objective lens (NA=1.46). Cells prepared and fixed as described above were covered with STORM buffer consisting of 25 mM HEPES, 5% Glycerol (v/v), Glucose oxidase at 25 µg/mL, Horse radish peroxidase 25 µg/mL and cysteamine 20 mM [27]. We ensured that out of focus drift is eliminated by setting the temperature of heating enclosure at 26^°^C and using the definite focus of the Elyra microscope. Alexa-647 flurophores were excited using 638 nm laser and emission was collected using long pass filter at 640 nm. Alexa-647 fluorophores were photo-converted from dark to excitable state, ~0.1mW of continuous 405 nm laser radiation and imaged with ≪1 mW of 488 nm light. Between 10,000-20,000 images were acquired per data set and single molecules emissions were fit using a 2D Gaussian distribution function in the algorithm provided in Zeiss’ Zen software. For image data acquisition we used a cooled, electron-multiplying charge-coupled device camera (iXon DU-897D; Andor) with an exposure time of 30 ms and an imaging area containing cells of interest. The zoom lens before the camera was set to 1.6X such that the pixel size was 0.1 µm. The data were acquired and pre-processed using the Zen (Zeiss) software customized for this microscope, while further cluster decomposition and statistics were done using custom built scripts in Matlab (The Mathworks Inc., Natick, MA). Pre-processing in Zen consisted of filtering out the features (molecules) detected with signal-to-noise parameter smaller than 6 and size smaller or equal to 9 pixels (mask) diameter which equaled 0.9 µm. No visible lateral (xy) or axial (z) drifts were observed in any data sets analyzed. No grouping of observation points was done, as we did not calibrate the blinking of Alexa-647 under imaging conditions in case they are uniformly randomly dispersed. Nevertheless, we do not claim that number of PIEZO1-GFP entities observed per cluster are in absolute values. Rather, we quantify the extent (area, perimeter) of clusters and quantify the relative change in observations, number of emitting events, per cluster in each condition. Pre-processed point pattern data was exported into a .txt file and loaded into Matlab for further quantification. Point pattern of observations, per cell, was segmented into clusters using the approach called DBSCAN [28] using custom built software in Matlab. Briefly, for each point detected, we calculated the local density within ~30 nm (average precision of detections in STORM using Elyra) radius. Next, we start with highest local density point and count all the molecules that lie within a circle of ~30 nm. Then, the same operation is repeated for every point that was within the circle, and so on until points at the boundary of a cluster have no further points included within their 30 nm radius. This is where all the points that passed the criteria are grouped into the first cluster. All the points from that cluster are removed from the original cluster data and stored. The operation is repeated until the next cluster gets isolated. This iterative process is repeated until the last cluster is defined and points included are stored. We search the objects that have at least 5 detections (points) per cluster, and any object with observations less than that is not considered a cluster. For each cluster we calculated the cluster area and perimeter using combination of Matlab functions (boundary, alphaShape, perimeter) and number of points per area (density) [29]. By defining the total cell surface, using the thresholded image from the green (GFP) channel, only the clusters within the cell boundaries were considered. This also allowed for counting number of PIEZO1-GFP clusters per unit cell area. Clusters smaller than the approximate size of a single PIEZO1 trimer (~200 nm^2^) were not considered for the statistics, as they could be the outcome of single molecule blinking, which we did not account for.

Clusters were excluded from the analysis based on the estimated number of molecules detected in each cluster. If the number of detected molecules in a given cluster was larger than the number of 200nm^2^ molecules (a conservative estimate of the area projected by a single PIEZO1 trimer) that could fit in that cluster area, then the cluster was rejected. This was formulated as Eq.1

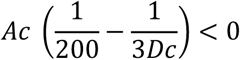

Where *Ac* is the measured *Cluster Area*, 200 is a conservative estimate of the area in nm^2^ projected by a single Piezo trimer, and *Dc is the Density of Molecular Detections/nm*^*2*^ detected in the same cluster, resulting in the following Area thresholding values Control (Ac > 269 nm^2^). This cutoff value was applied to all experimental conditions.

### Live cell TIRF imaging and k-space Image Correlation Spectroscopy (kICS) confinement analysis

PIEZO1-GFP diffusion inside the plasma membrane of stably transfected HEK293T cells was imaged by a TIRF Elyra (Zeiss) microscope using 100X/1.46 lens at an imaging rate of 29 Hz. The microscope stage was equipped with an enclosed heating module (37 °C). To quantify the diffusion of PIEZO1-GFP clusters and its confinement by cholesterol domains, we applied the k-space Image Correlation Spectroscopy (kICS) confinement analysis, as previously reported [29]. By assuming the isotropic diffusion of fluorescent particles, we circularly averaged the correlation function at each temporal k^2^ and temporal lags ‘tau’, as shown in examples of SI Fig. 7A-C. Subsequently, we fitted the resulting correlation function (SI Fig. 7D) to extract the diffusion (SI Fig. 7E) of PIEZO1-GFP outside of cholesterol clusters, and the lateral unbinding rate (SI Fig. 7E) of PIEZO1-GFP from cholesterol rich domains, as described in equation 2 of reference [30]. The kICS correlation functions from late temporal lags and for large spatial frequencies was fitted using equation 9 of reference [30].

### Live PIEZO1-GFP and CtxB-Alexa-555 imaging and co-localization by Spatio-Temporal Image Cross Correlation Spectroscopy (STICCS)

Live PIEZO1-GFP stable cells were labelled with CtxB-Alexa-555 and imaged on a ZEISS Elyra microscope in TIRF mode, as described above.For two color imaging on a single EMCCD camera, we used a dual band dichroic filter cube for 488/561 nm excitation, where an AOTF laser controlling module was used to separately excite GFP (Piezo1) or Alexa-555 (CtxB) at any given frame. The resulting frame time was increased to about 240 ms, while exposure time was set at 33 ms. We quantified the degree of co-localization between PIEZO1-GFP and the cholesterol domain marker, CtxB-Alexa-555, and their diffusion rate, by Spatio-Temporal Cross-Correlation Spectroscopy [31, 32]. For each image series, in two channels, a rectangular area surrounding an individual cell was defined. The immobile (DC) population from each cell was filtered by Fourier filtering, from each channel. STICS correlation function was calculated for each channel and for cross-correlation, as described in [31, 32]. The amplitude of auto- and cross-correlation functions were used to calculate the density of co-localized entities using the following equation: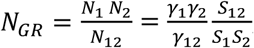, where γ_1_, γ_2_ and γ_12_ are geometrical factors of the point spread function (PSF) in each channel and S_1_, S_2_and S_12_ are slopes of linear fit between the amplitudes and the inverse squared widths of correlation function, as measured in each temporal lag, τ [32]. In order to characterize noisier STICCS correlation functions, from short temporal sequence data sets and obtain squared widths, σ^2^, of correlation functions, we employed the 2D moment analysis as previously used in equation 10 of ref [33]. For amplitudes, we used the assumption that the integral under the curve of a 2D Gaussian will be equal to *π*(*amplitude*)σ^2^ and calculating the numerical integral at each temporal lag. The PSF gamma factors can be calculated using 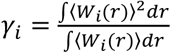 for each channel i separately and 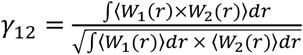 for cross-correlation. To measure gamma factors, we imaged a z-stack of 0.1 μm sized tetraspeck (INVITROGEN, etc) nanoparticles and calculated the above integrals numerically. The resulting ratios between gammas were 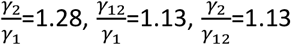, for the imaging configuration described above. The number of PIEZO1 entities at each condition was obtained by applying the analysis described above to the image data for the PIEZO1 imaging channel (488 nm) and using equation 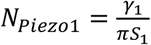. To characterize the diffusion of co-localized entities of PIEZO1 and CtxB, we used linearly fit squared widths, s^2^ of STICCS correlation functions vs temporal lag τ, and the resulting slopes are equal to 4Dτ [31, 32].

To furthermore investigate the effect of MBCD on PIEZO1 co-localization with CtxB, we performed experiments where the same cells were imaged prior and post treatment. We imaged cells pre-treatment as described above while recording their positions and added drugs to the dish ensuring that the dish could not move in order to preserve the saved cells coordinates. Live cell imaging acquisition began 20 minutes after treatment was applied.

### Imaging membrane order with Laurdan and ICCS analysis

Cells plated on dishes were labelled with 1:200 dilution of 5 mM Laurdan (D250, Molecular Probes), incubated at 37 °C for 30 min, washed with warm PBS and fixed, permeabilized and then the PIEZO1-GFP was labeled with Anti-GFP-Alexa-647.

Laurdan data was acquired on a Zeiss LSM 880 microscope with excitation from a mode-locked two-photon Insight DeepSea operating at 80 MHz repetition rate and a wavelength of 780 nm. Excitation and photon collection were performed by Plan-Apochromat 63x 1.4 Oil objective (Zeiss-GmbH, Germany). Laurdan data was acquired in 2-channels with wavelength range 400-460 and 470-530 nm respectively. Image Cross-Correlation Spectroscopy (ICCS) was applied to quantify the level of co-localization between PIEZO1 and ordered/disordered phases of Laurdan [34]. Each image pair was pre-processed to minimize the artefacts that can be due to the channels misalignment, noise and cell edges [34]. First, a correction matrix for shift of far red (Alexa 647) images to align with Laurdan images, was calculated using Matlab function imregtform. The corrected images were then noise filtered using a 2D Gaussian filter with 1 pixels sigma. Then the minimum intensity from images, in each channel was subtracted and a final image was normalized by its maximum. Finally, the resulting images were used to define a mask, from thresholding above the average intensity, and all the pixels not belonging to the mask (equal to 0) were set to have the value of average intensity of pixels within the mask (equal to 1). This ensures that pixels not belonging to cell areas, are not contributing to the fluctuations and inherent correlation functions in ICCS [34]. We extract the number density of particles in red (PIEZO1), green (either ordered or disordered Laurdan phase) and cross-correlation channels.

## RESULTS

### Electrophysiological effects of membrane cholesterol manipulation

We measured the effects of cholesterol depletion, supplementation and dynasore (a compound known to disrupt cholesterol rich domains) on PIEZO1-GFP channels stably expressed in HEK293T cells. These cells express exceedingly low levels of STOML3 [25, 26]. We observed a significant modulation of channel sensitivity and kinetics when either cholesterol was removed by MBCD or dynasore was added as compared to control cells (Fig. 1A-D). High doses (100 μg/ml) of water-soluble cholesterol (Fig.1B) did not affect PIEZO1-GFP gating. However, PIEZO1-GFP pressure sensitivity was significantly decreased after treatment with 5 mM MBCD (Figs. 1E-F) and this effect was accompanied by a substantial increase in latency of the response, both at the level of the first opening (SI Fig.1) and onset of the maximal current (Fig. 1G). None of the compounds altered the average maximal PIEZO1-GFP current (Fig. 1H). Both MBCD and dynasore caused similar delays in PIEZO1-GFP activation but only MBCD caused a right-ward shift in the midpoint activation pressure of the channel by ~40%. Interestingly, both MBCD and dynasore treatment largely abolished channel inactivation. This was quantified using normalized steady state current (Fig 1I). Similar effects of MBCD were observed on endogenous mouse PIEZO1 channels in N2A cells (SI Fig.2).

**Figure 1.**
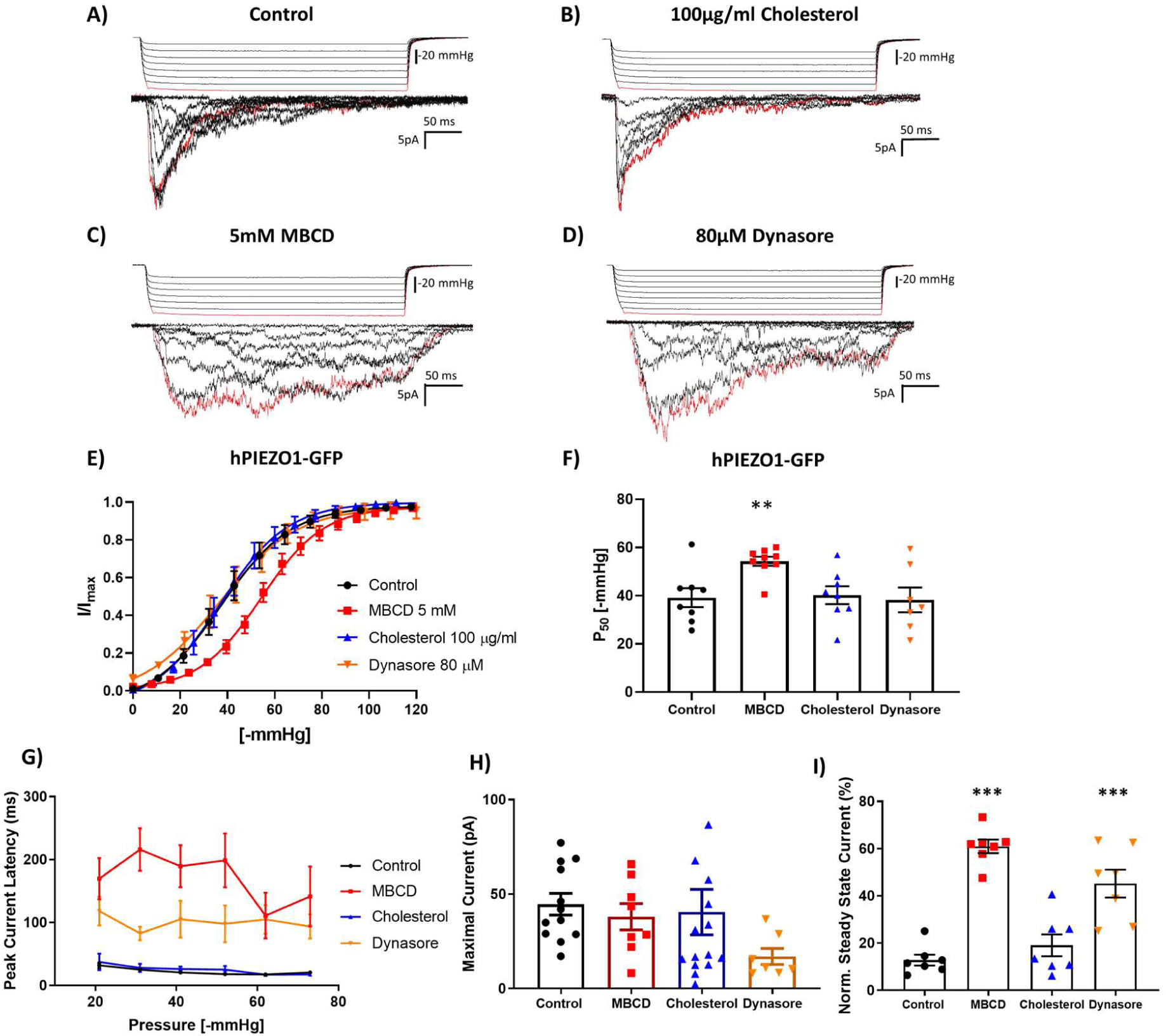
Modulation of PIEZO1-GFP activity in cell-attached patches of HEK293T cells in response to manipulation of cholesterol domains in the plasma membrane. (A-D) Example recordings from cell attached patches in response to the application of negative pressure pulses of HEK293T cells stably expressing PIEZO1-GFP at +65mV pipette voltage. Currents shown are from (A) control cells incubated in serum free media for 30 minutes, (B) cells treated with 100 μg/ml water-soluble cholesterol, (C) cells incubated in serum free media supplemented with 5mM MBCD or (D) supplemented with 80μM dynasore (The first pressure pulse and its associated current are highlighted in red). (E) Pressure response curves comparing the effects of each compound on PIEZO1-GFP pressure sensitivity. Peak current values (I) are normalized to the maximal current (I_max_) within the same recording and fitted with a Boltzmann distribution curve. (F) Midpoint activation pressures (P_50_) producing 50% of the maximal current response from each Boltzmann curve analysed in panel (E) (One-Way Anova ** P < 0.005; n.s. – not significant, n= 7-10 cells). (G) Latency time in ms of the PIEZO1-GFP response measured between the start of negative pressure pulse and the onset of the maximal current within the same recording (n=3, where each is the average of at least 3 consecutive pressure protocols). Average latencies are displayed for pulse pressures between −20 and −70 mmHg. (H) Average maximal current from cell-attached patches generated from each cell in all conditions. (I) Quantification of the mean steady state current produced by each cell. The average current measured over the last 100ms of the pressure stimulus was normalized to the maximal current within the same cell. (n = 7 cells per condition. The steady state current for each cell is the average mean steady statute current from a minimum of 3 pressure pulses between −40 and −60 mmHg. All data represents mean ± S.E.M.; Statistical analysis was performed using One-Way ANOVA (** P < 0.005, *** P < 0.0005).

### Cholesterol removal exacerbates the slow-inactivating phenotype of PIEZO1 R2456H

The inactivation of PIEZO1 is currently described as an intrinsic process in the channel gating cycle, which can be altered by introducing mutations at specific sites in the protein [23, 35-37]. The R2456H mutant, a xerocytosis-causing variant of human PIEZO1, displays a significant loss of inactivation with respect to wild-type (WT) PIEZO1 (Fig.2A), as previously reported [6]. We tested whether the effects of MBCD treatment on the channel inactivation were specific to the WT channel or could effect the R2456H channel as well (Fig.2A-B). The degree of inactivation was quantified using normalized steady state current (Fig.2A-C). Interestingly, the MBCD treatment appeared to slow inactivation of the R2456H PIEZO1 (Fig.2B-C) increasing the steady state current while also increasing the latency of the response (FIG.2D, SI Fig.1) when the mutant was transfected into PIEZO1 KO HEK293T cells. MBCD did not change the sensitivity of R2456H to applied pressure (Fig.2E-F).

**Figure 2.**
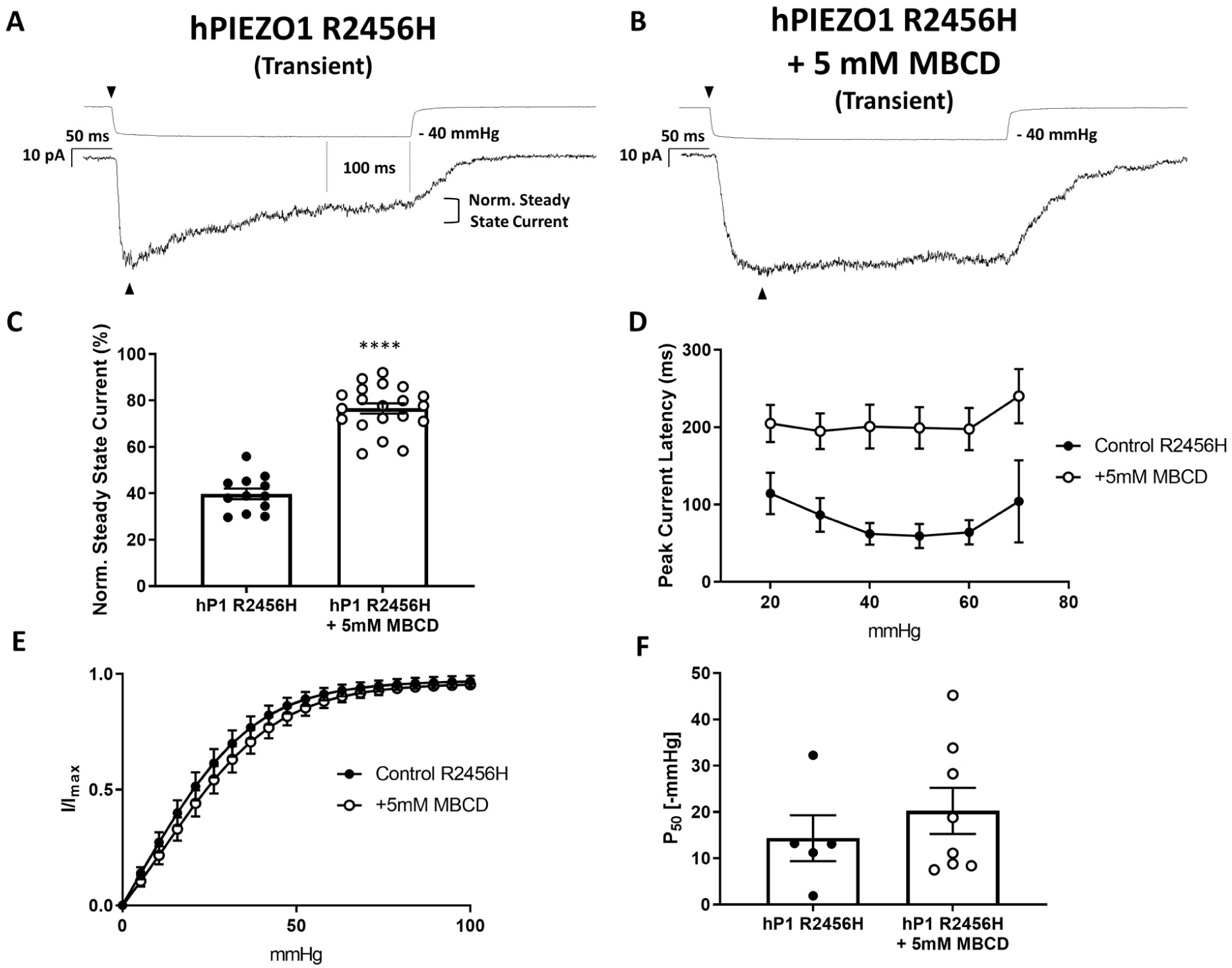
The slow-inactivation phenotype of the human PIEZO1 mutant R2456H is exacerbated by the removal of plasma membrane cholesterol via MBCD. The R2456H mutation was introduced in the human PIEZO1-1591-GFP clone and henceforth referred to as hPIEZO1 R2456H. (A) Example trace of a cell-attached recording from transiently over-expressed human hPIEZO1 R2456H mutant. (B) Example trace of a cell-attached recording from transiently over-expressed human hP1 R2456H treated with 5mM MBCD treatment for 30 minutes in serum free media. (C) Histogram quantifying the impact of 5mM MBCD treatment on the normalized steady state current. [hPIEZO1 R2456H (n = 12); hPIEZO1 R2456H + 5 mM MBCD (n = 20)]. (D) This effect was accompanied by a longer latency to activation of the peak current at all pressures after treatment with MBCD [Control (n = 6), MBCD (n = 8)]. (E) Pressure response curve of hPIEZO1 R2456H and MBCD-treated hPIEZO1 R2456H showing Boltzmann curve fits. [hPIEZO1 R2456H (n = 5), hPIEZO1 R2456H + MBCD (n = 8)]. F) Quantification of P_50_ values obtained from Boltzmann fits analysed in E. Statistical analysis was performed using One-Way ANOVA (**** P < 0.0001).(n = 1 represents the average of minimum 3 recordings from the same cell). Error bars in all graphs represent Standard Error of the Mean (S.E.M).

Interestingly, cholesterol removal via MBCD did not significantly change the midpoint of activation of the transiently transfected WT PIEZO1 to applied pressure either (SI Fig.3A-B). We hypothesized that the shift in pressure sensitivity observed in our stable cell line after MBCD treatment was dependent on the mode of the channel expression (i.e. stable vs transient). Apparently, PIEZO-GFP from the stable cell line would be more tightly regulated in expression levels and localization over several generations, while transiently overexpressed channels would bypass such regulation and incorporate in the plasma membrane at random. The transiently expressed R2456H mutant displayed a higher sensitivity to pressure (Fig.2F) compared to the transiently expressed WT PIEZO1 (SI Fig.3B) and stably expressed PIEZO1-GFP (Fig.1 F). The P_50_ values and slopes of Boltzmann fits are summarized in Table 1. Only the stable cell line and the PIEZO1 native to N2A cells (SI Fig.2), appeared to right shift their sensitivity after MBCD treatment. All the PIEZO1 variants studied (PIEZO1-GFP, transiently transfected WT, transiently transfected R2456H mutant and N2A native) displayed slower inactivation after cholesterol removal. These results suggest that membrane cholesterol, like other lipids [20], contributes to PIEZO1 inactivation.

**TABLE 1.** Summary of average P_50_ values and slopes derived from Boltzmann fits of I/I_max_ versus Pressure plots of all PIEZO1 variants and treatments performed in this study. Gating free energy calculations were estimated following eq. 7 (ΔG/k_B_≈ P_50_ · α) in [38]. Statistical difference between ΔG detected for PIEZO1-GFP vs. 25 μM DHA [24 hr] (* P = 0.0153, Student’s T-test). All values are reported as mean with S.E.M.

| | $P_{50}$ [-mmHg] | Slope of Boltzmann<br>$\alpha^{-1}$ [mmHg<sup>-1</sup>] | $\Delta G$ ( $k_B T$ ) |
| --- | --- | --- | --- |
| hPIEZO1-GFP | 36.4 ± 3 | 14 ± 2.5 | 3.5 ± 0.4 |
| + 5mM MBCD | 54.2 ± 1.1 | 13.3 ± 1 | 5.3 ± 1.2 |
| + 100 µg/ml Cholesterol | 36.8 ± 2.5 | 14.4 ± 2 | 3.5 ± 0.2 |
| + 80 µM Dynasore | 36.7 ± 4.8 | 16.3 ± 3.9 | 3.7 ± 1 |
| + 25 µM DHA [24 hr] | 16 ± 2.8 | 12.3 ± 1.4 | 1.6 ± 0.4 |
| + 25 µM DHA [48 hr] | 27 ± 1.8 | 10.3 ± 1.4 | 3.6 ± 0.8 |
| hPIEZO1 WT | 39.6 ± 2.1 | 11.6 ± 2 | 4.4 ± 0.5 |
| + 5 mM MBCD | 42.2 ± 2.6 | 14.3 ± 2.1 | 3.7 ± 0.6 |
| hPIEZO1 R2456H | 13.3 ± 7 | 15.1 ± 3 | 1.3 ± 0.6 |
| + 5 mM MBCD | 16.4 ± 5.6 | 14.1 ± 2.8 | 1.9 ± 0.5 |
| N2A mPIEZO1 | 17.8 ± 2.2 | 6.6 ± 1.6 | 4.7 ± 0.3 |
| + 5 mM MBCD | 22.5 ± 1.8 | 8.4 ± 1.7 | 4 ± 0.7 |

### Cholesterol depletion increases PIEZO1-GFP diffusion in live cells

To study the dynamics of PIEZO1-GFP in a live cell we performed spatio-temporal kICS analysis of pixels’ fluorescence fluctuations in the basal membrane of our stable cell line (Fig. 3A). The PIEZO1-GFP fluorescent signal in TIRF mode was detected as distinct puncta of varying sizes. The smaller fluorescent entities diffusing away from large puncta were defined as clusters while puncta larger than point spread function (~200 nm radius) were defined as group of clusters (Fig.3B). The term cluster was used to indicate a diffraction-limited fluorescent entity composed by an unknown amount of PIEZO1-GFP proteins. PIEZO1-GFP appears to experience a limited diffusion, schematically described in the cartoon as the motion from A to B (Fig. 3B), due to its clustering and heterogeneity of the plasma membrane. Prior to the the addition of MBCD, the channel’s average diffusion rate was ~0.005 μm^2^/s (Fig. 3C). Cholesterol removal by MBCD doubled the diffusion rate while addition of 50 µg/ml of cholesterol had no effect. Also, kICS analysis measures the lateral unbinding rate from the PIEZO1-GFP clusters, k_unbind. This measurement is defined as the rate at which fluorescent entities, particles or clusters, move away from locally confined areas or docking areas. This parameter can also be interpreted as rate at which a measured protein escapes a local confining environment. MBCD treated cells caused PIEZO1-GFP molecules to dissociate at a higher rate from associated local confinement while cholesterol addition did not affect this affinity (Fig.3D). This implies that MBCD treatments breaks the underlying local cholesterol scaffold by removing the cholesterol and changing the affinity of PIEZO1-GFP to its local environment while cholesterol addition does not change the confinement barrier, but only the number of barriers. Perturbing the actin (CytochalasinD, Jasplakinolide) or the microtubule cytoskeleton (Colchicine) did not affect the local lateral unbinding of the PIEZO1-GFP from clusters (SI Fig.4A). Interestingly, among these compounds, only jasplakinolide appeared to reduce the effective diffusion in the basal membrane, consistent with its effect on the organization of the cortical actin meshwork (SI Fig.4B).

**Figure 3.**
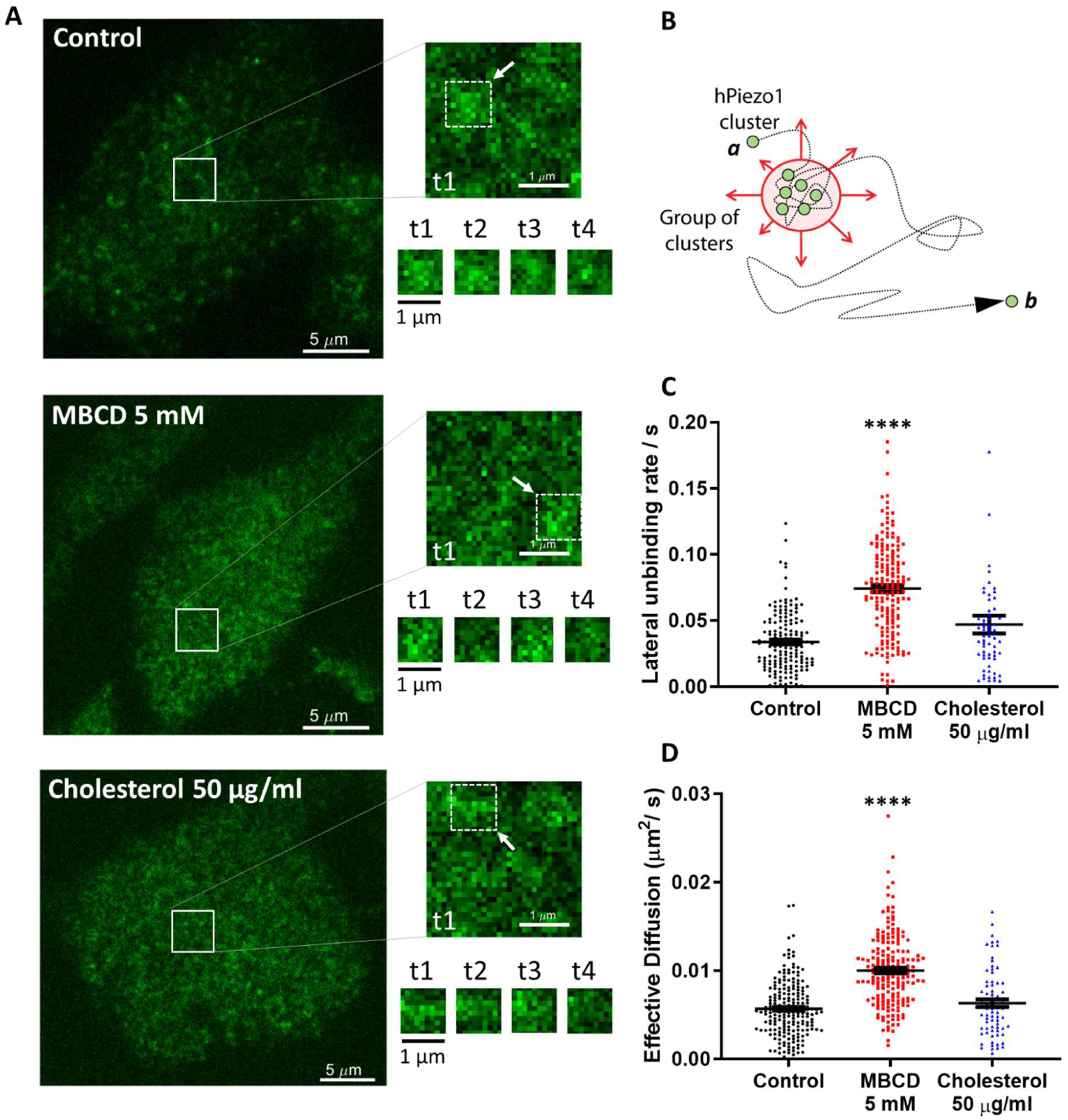
Cholesterol removal alters PIEZO1-GFP dynamics in situ. (A) Left panels, TIRF images are the initial frames acquired during live cell imaging. The right panel shows a representative time course of PIEZO1-GFP diffusion dynamics within the highlighted square area. This is shown next to each of the conditions tested. The four time points (t1 to t4) represent the time course of the fluctuations in fluorescence intensity within the 1 µm^2^ portion of the image indicated by the white arrow. (B) Cartoon describing the limited diffusion experienced by PIEZO1-GFP channels in the plasma membrane. (C) Quantification of lateral unbinding rate (s^2^) of PIEZO1-GFP entities in control, 5mM MBCD treated and 50 μg/ml cholesterol treated cells. (D) Effective diuffusion (μm^2^/s) of PIEZO1-GFP entities in control, 5mM MBCD treated and 50 μg/ml cholesterol treated cells. Both lateral unbinding rates and effective diffusion are increased significantly by cholesterol depletion via MBCD. (Data represents mean ± S.E.M. analyzed with One-way ANOVA *P < 0.05, **** P < 0.00005).

### Nanoscale organization of PIEZO1-GFP clusters and cluster dynamics

To confirm our observations on PIEZO1-GFP clustering dynamics, we performed Stochastical Optical Reconstruction Microscopy (STORM) in TIRF mode. By applying STORM we were able to detect the boundaries of each fluorescent entity with nanometric precision and therefore describe the biophysical characteristics of large and small PIEZO1-GFP clusters.

Our stable cell line carrying a PIEZO1-GFP fused construct permitted us to visualize stochastic blinking events originating from an Alexa647-labelled anti-GFP antibody. This allowed us to avoid issues with the specificity of commercially available primary PIEZO1 antibodies. Figure 4A shows the PIEZO1-GFP stable cell line and the resolution improvement due to STORM analysis. The insets in the figure focus on subcellular areas where the GFP signal (in green) was superimposed with the STORM signal (in red) inferred from the optical reconstruction of Alexa-647 blinking events. Software cluster decomposition and classification allowed us to characterize PIEZO1-GFP clusters by defining their Area (Fig. 4B), perimeter length (Fig. 4C) and cluster density/μm^2^ (Fig. 4D).

**Figure 4.**
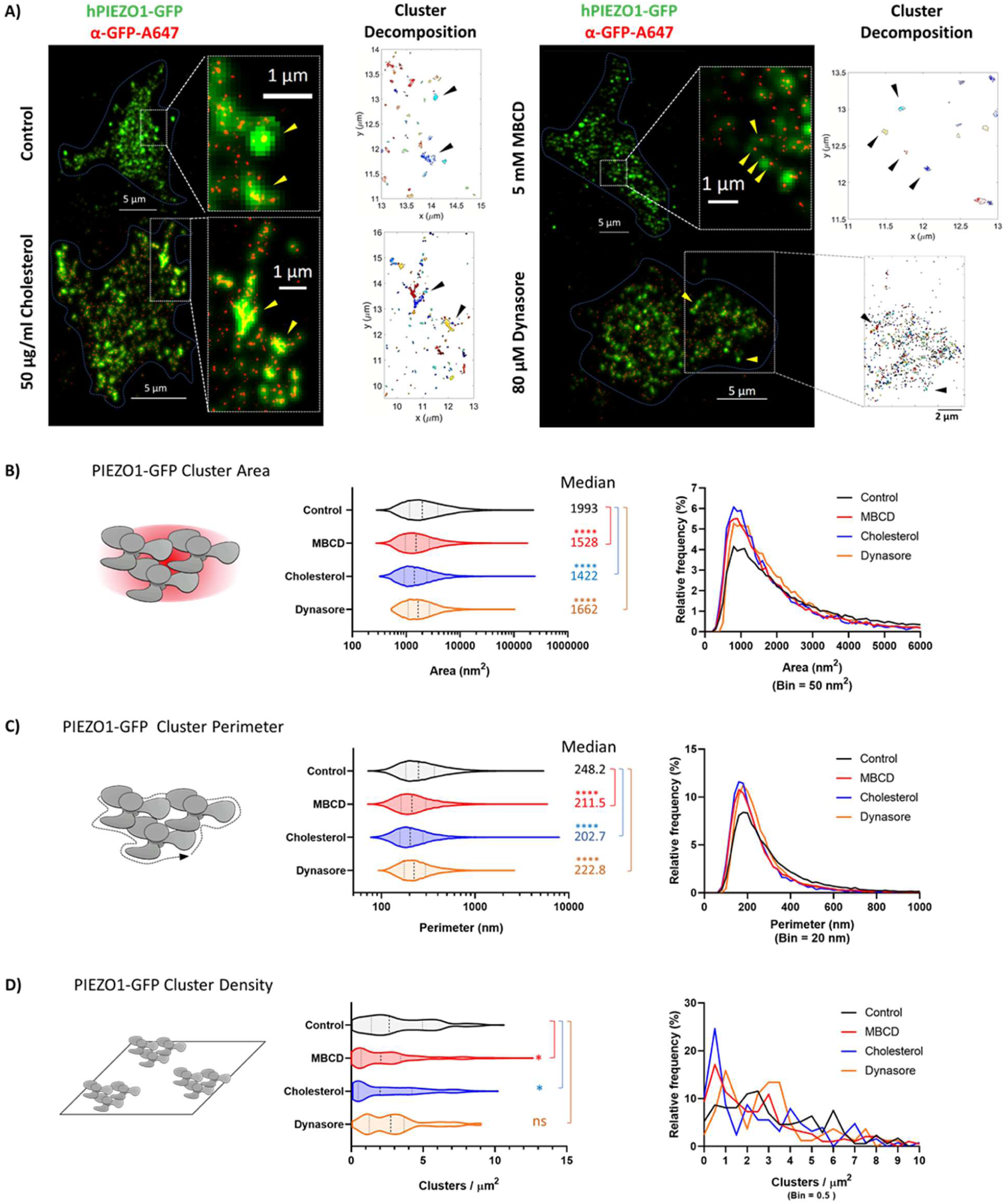
Modulation of PIEZO1-GFP clusters determined by STORM super-resolution microscopy. (A) TIRF microscopy of PIEZO1-GFP cells immunolabelled with an anti-GFP antibody conjugated to Alexa647. Images of control cells and treated cells (Cell boundary highlighted by a dashed blue line. Scale bar represents 5 μm) show the PIEZO1-GFP signal (green) superimposed on the STORM signal from Alexa647 (red)-see insets. The physical properties of the clusters shown in the insets were analysed by Cluster Decomposition and are shown in the adjacent cluster maps. (B) Left hand panel illustrates quantification of PIEZO1-GFP cluster area as violin plots in control and treatment conditions. Right hand panel shows the relative frequency of cluster areas in nm^2^. (C) Left hand panel illustrates quantification of PIEZO1-GFP cluster perimeters as violin plots in control and treatment conditions. Right hand panel shows the relative frequency of cluster perimeters in nm [For cluster area and perimeter: Control PIEZO1-GFP (n = 34741), MBCD (n = 19159), Cholesterol (n = 12097), Dynasore (n = 19982). n=1 represents a single PIEZO1-GFP cluster]. (D) Left hand panel illustrates quantification of PIEZO1-GFP cluster density as violin plots in control and treatment conditions. Right hand panel shows the relative frequency of cluster density in clusters per μm^2^ [Control PIEZO1-GFP (n = 173), MBCD (n = 193), Cholesterol (n = 126), Dynasore (n = 82). N = 1 represents a single cell]. All violin plots are marked with median (thick dashed line) flanked by 25- and 75-percentile (thin dashed lines). The median value of each plot is reported on the right hand-side of each graph. (Statistical analysis: One-Way Anova, * P < 0.05, **** P < 0.00005, ns = not significant).

In control experiments PIEZO1-GFP clusters appeared to distribute non-uniformly around the cell’s basal membrane and to pack into a variety of cluster areas and perimeters. Cluster areas ranged from few hundred nm^2^ to several thousand nm^2^. Less than 10% of total clusters in relative frequency histograms, appeared to be ~1000-2000 nm^2^ in area (Fig. 4B) and ~200-300 nm in perimeter (Fig. 4C). All tested conditions appeared to increase the relative frequency of this fraction of clusters, indicative of breakdown of larger order clusters into smaller ones. Despite the statistical difference detected between the median values for cluster sizes between conditions, the distribution of these cluster populations largely overlapped. In addition the stably expressing HEK293T cells displayed variable levels of cluster densities in their basal membrane (Fig.4D), and we identified 3 major populations: low-PIEZO1-GFP cells (~2-3 clusters / µm^2^), medium-PIEZO1-GFP cells (~6 clusters / µm^2^) and high-density PIEZO1-GFP cells (~8-9 clusters / µm^2^). As suggested by the size histograms, each treatment appeared to impact the density of PIEZO1-GFP clusters in the basal membrane by increasing the number of cells displaying reduced cluster densities. Out of all tested conditions, dynasore appeared to uniformly reduce the cluster density in all 3 observed groups. MBCD and cholesterol treatments, on the other hand, greatly reduced the fraction of cells displaying high densities of PIEZO1-GFP, enriching the fraction of cells displaying low density PIEZO1-GFP clusters.

These results suggest that PIEZO1-GFP assembles into clusters in the plasma membrane of cells. Cholesterol levels in the plasma membrane appear to regulate the degree of clustering by mostly affecting the density of clusters in the basal membrane.

### Colocalization of PIEZO1-GFP and raft marker cholera toxin subunit B (CtxB)

To visualize association of PIEZO1-GFP and cholesterol-rich domains in stably expressing HEK293T cells, we used fluorescently labelled (Alexa555) Cholera Toxin subunit B (CtxB), a well-known marker of lipid rafts [39]. Because of the heterogeneous and dynamic behavior of PIEZO1-GFP entities in the membrane, we collected imaging data as time-lapse series in two channels and examined the degree of cross-correlation between GFP- and Alexa555-tagged entities using STICCS [see Methods]. This approach allows for quantitation of density of fluorescent entities per µm^2^ (amplitudes of correlation functions) and to calculate the number of overlapping (co-localized) entities in a series as well as their dynamics. In control experiments, PIEZO1-GFP appeared as punctate entities consistent with channel clustering that often co-localized with CtxB-A555 (Fig. 5, top row). Quantification of PIEZO1-GFP/CtxB co-localization revealed that PIEZO1-GFP co-localizes with CtxB at about 40% (SI Fig.5A). These co-localized entities dissociated upon addition of MBCD (Fig. 5A, second row). This resulted in a larger fraction of smaller PIEZO1-GFP entities detected in the membrane (Fig. 5B) as well as an increase in colocalized entities (Fig.5C). The density of fluorescent CtxB didn’t change across conditions indicating that GM1 ganglioside levels weren’t affected (SI Fig.5B).

As previously observed (Fig. 3D), the MBCD-treated PIEZO1-GFP entities also appeared to diffuse faster in the basal membrane than in control consistent with the idea that cholesterol-rich domains spatially confine PIEZO1-GFP. Interestingly, the diffusion speed of co-localized PIEZO1-GFP/CtxB entities was faster after MBCD treatment when compared to the diffusion speed of co-localized entities in the control condition (Fig.5D). The MBCD treatment approximatively doubled the diffusion speed of both individual CtxB entities (SI Fig.5C) and of the PIEZO1-GFP/CtxB colocalized entities (Fig.5D). The effects of MBCD treatment were also observed in time-course experiments at the single cell level (SI Fig.6). Dynasore- and cholesterol-treated cells showed no significant difference in either PIEZO1-GFP cluster numbers (Fig. 5B) or their co-localization with CtxB (Fig.5C) when compared to control. In addition, CtxB did not significantly affect the diffusive dynamics of PIEZO1-GFP.

**Figure 5.**
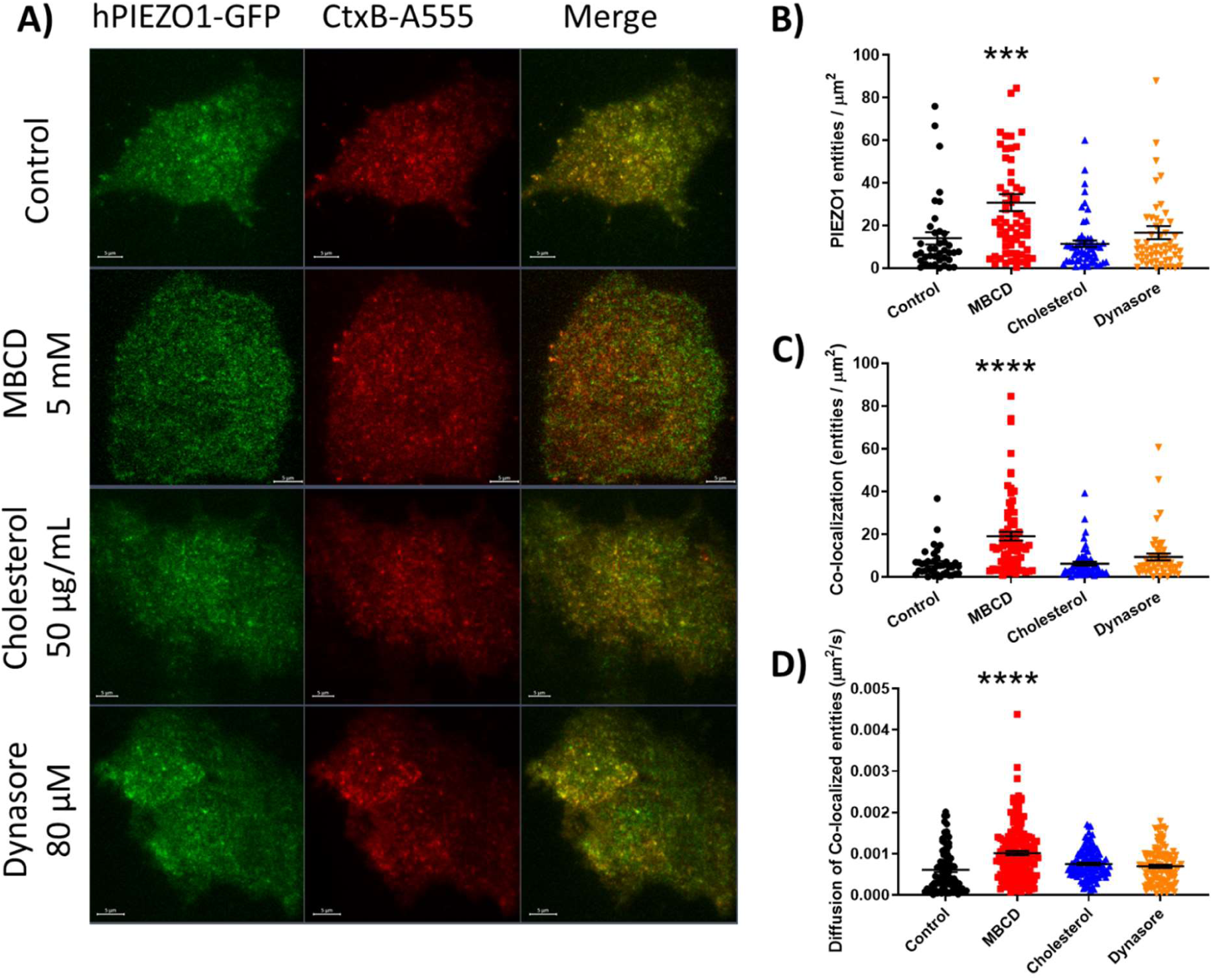
Co-localization of PIEZO1-GFP and CtxB raft marker using TIRF. (A)Total Internal Reflection Fluorescence (TIRF) microscopy of cholesterol-rich domains labeled with CtxB-A555 (red) on the basal membrane of PIEZO1-GFP expressing HEK293T cells (GFP fluorescence in green). (B) Average number of PIEZO1-GFP entities per μm^2^ detected after 30 min treatments. (C) Average density of co-localized entities per μm^2^ detected after 30 min treatments. D) Effective diffusion rate of co-localized PIEZO1-GFP/CtxB-A555 at the micron scale, before and after treatments. (One-Way ANOVA, *** P < 0.0005, **** P < 0.0001; error bars represent S.E.M). Scale bars represent 5 μm.

Collectively, these results suggest that PIEZO1-GFP associates into larger clusters in a cholesterol-dependent manner. Disruption of cholesterol organization in the membrane facilitates diffusion of PIEZO1-GFP suggesting a confinement by cholesterol-rich fences.

### Colocalization of PIEZO1-GFP and the membrane order probe Laurdan

Using 2-photon microscopy (Ex: 780 nm) we were able to distinguish the fluorescent signals emitted from Laurdan associated with lipid-ordered membrane domains (em: 400-460 nm) and lipid-disordered ones (em: 470-530 nm) and assessed the degree of co-localization between each lipid phase and PIEZO1-GFP using Image Cross Correlation Spectroscopy (ICCS). As previously observed, the density of PIEZO1-GFP fluorescent entities appeared to increase upon cholesterol domain disruption after MBCD and dynasore treatments (Fig. 6A). The density of ‘ordered’ Laurdan (fraction of Laurdan associated with ordered lipids) was increased upon cholesterol enrichment (Fig. 6B). Conversely, Laurdan phase distribution appeared to be sensitive to cholesterol disruption, displaying a significant increase in detectable ‘disordered’ Laurdan entities (fraction of Laurdan associated with disordered lipids) after exposure to dynasore (Fig.6C), but surprisingly not after MBCD treatment. In the control condition PIEZO1-GFP appeared to co-localize more often with disordered Laurdan entities (Fig.6D, control = 9.7 co-localized entities / µm^2^) than their ordered counterpart (Fig.6E, control = 18.5 co-localized entities / µm^2^). The dynasore treatment increased the average density of PIEZO1-GFP co-localized with both the ordered (Fig.6D, Dynasore = 19.1 co-localized entities / µm^2^) and disordered (Fig.6E, control = 52.8 co-localized entities / µm^2^) membrane regions detected by Laurdan. Cholesterol enrichment did not affect the density of PIEZO1-GFP colocalized with ordered lipids, despite increasing the fraction of ordered Laurdan. These results suggest that under control conditions PIEZO1-GFP is preferentially localized in lipid-disordered membrane domains. Dynasore treatment enhances phase separation of Laurdan by increasing the density of disordered membrane domains, causing a greater fraction of PIEZO1-GFP to associate with disordered lipids. The MBCD treatment didn’t appear to significantly increase the fraction of disordered membrane domains while increasing the density of PIEZO1-GFP entities.

**Figure 6.**
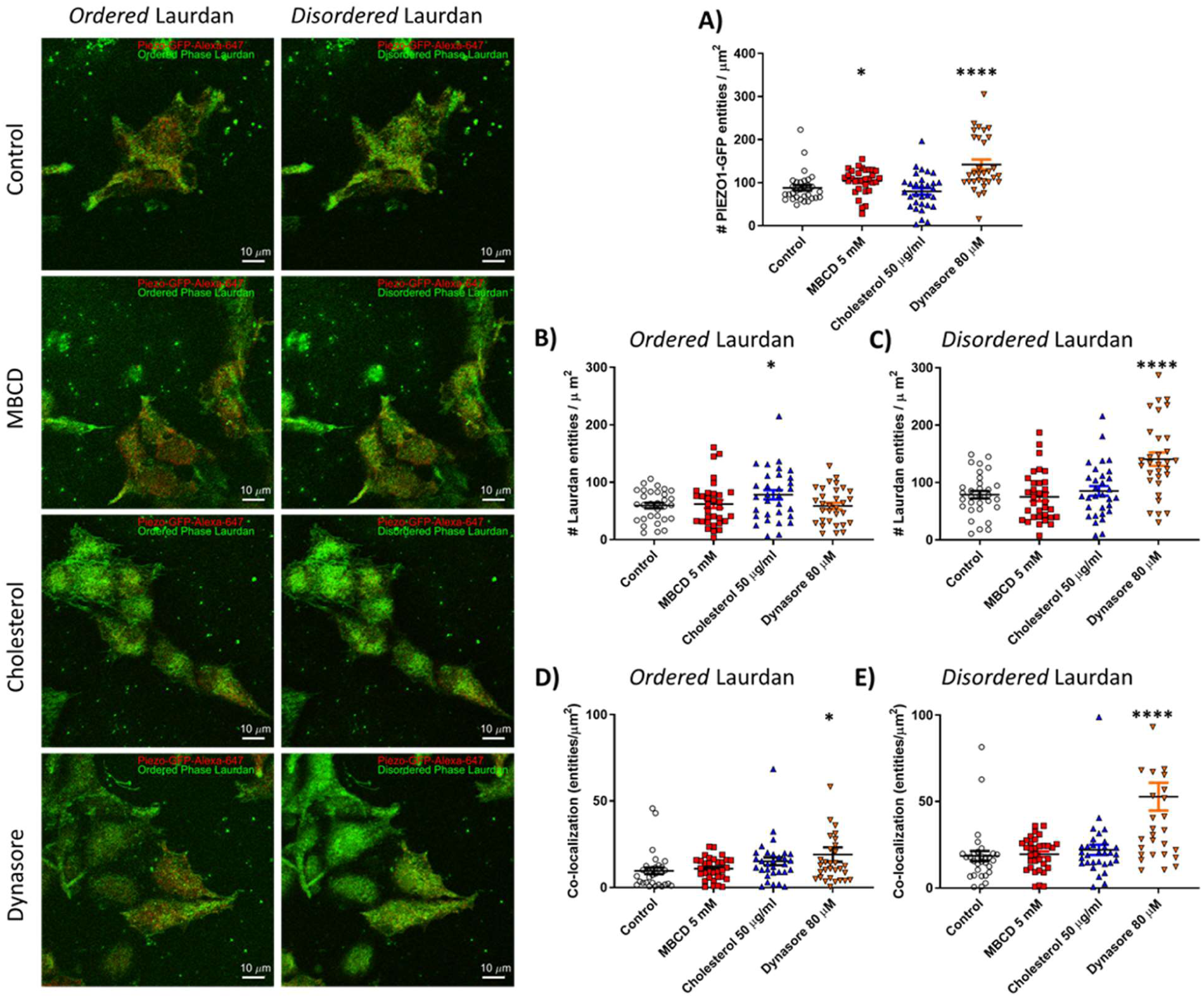
Co-localization between PIEZO1-GFP and Laurdan phases. Left) fluorescence microscopy of hP1-1591-CL cells labelled with Laurdan. PIEZO1-GFP (red) is tested for colocalization with the Laurdan dye (green). The images on the left column display the signal emitted by ordered-phase Laurdan (em: 400-460 nm) and the right column shows the signal emitted by disordered-phase Laurdan (em: 470-530 nm). A) Density of PIEZO1-GFP entities per µm^2^ detected under various conditions. B) Density of ordered Laurdan entities. C) Density of disordered Laurdan entities. D) Density of colocalized PIEZO-GFP/Ordered Laurdan entities as determined by Image Correlation Spectroscopy (ICS). E) Density of colocalized PIEZO-GFP/Disordered Laurdan entities. N = 1 represents data obtained from a single cell. Control PIEZO1-GFP (n = 31 cells), 5 mM MBCD (n = 32 cells), 50 μg/ml cholesterol (n = 31 cells), 80 μM dynasore (n = 31 cells). Statistical analysis was performed using One-Way ANOVA, * P < 0.05, **** P < 0.0001; error bars represent S.E.M.

### Sensitization of PIEZO1-GFP current by DHA supplementation

Given that we saw PIEZO1 preferentially localized in regions with a higher disorder we next sought to see how poly-unstaurated fatty acids affected PIEZO1-GFP. Poly-unsaturated fatty acids are one likely candidate to comprise these regions of disorder. The fatty acid DHA is an important dietary poly-unsaturated fatty acid (PUFA) which is thought to enhance separation of lipid phases in vitro and confer fluidity to disordered lipid phases [40]. The fatty acid becomes incorporated into membrane phospholipids when supplied long-term in the cell culture medium [41]. Under standard pressure protocol conditions, DHA-treated cells displayed a mixed phenotype of inactivating currents at low pressure regimes (blue trace) and non-inactivating currents at higher pressure regimes (red trace) as exemplified in Fig.7A. Long-term incubation of stably expressing PIEZO1-GFP cells with 25 µM DHA for 24 hr and 48 hr left-shifted the dose-response curve of PIEZO1-GFP in cell attached patches (Fig.7B, 7C). The DHA supplementation increased the latency of the maximal current (Fig.7D) and of the first current event (SI Fig.1). Quantification of channel inactivation over the pressure range tested revealed a significant loss of inactivation when the channel was stimulated with pressure pulses greater than −30 mmHg (Fig.7E). These results indicate that DHA can sensitize the PIEZO1-GFP response to applied pressure (Table 1) while simultaneously delaying the onset of both the first event and the peak-current. Furthermore, sensitization of the channel can cause it to adopt a non-inactivating gating mode when over-stimulated.

**Figure 7.**
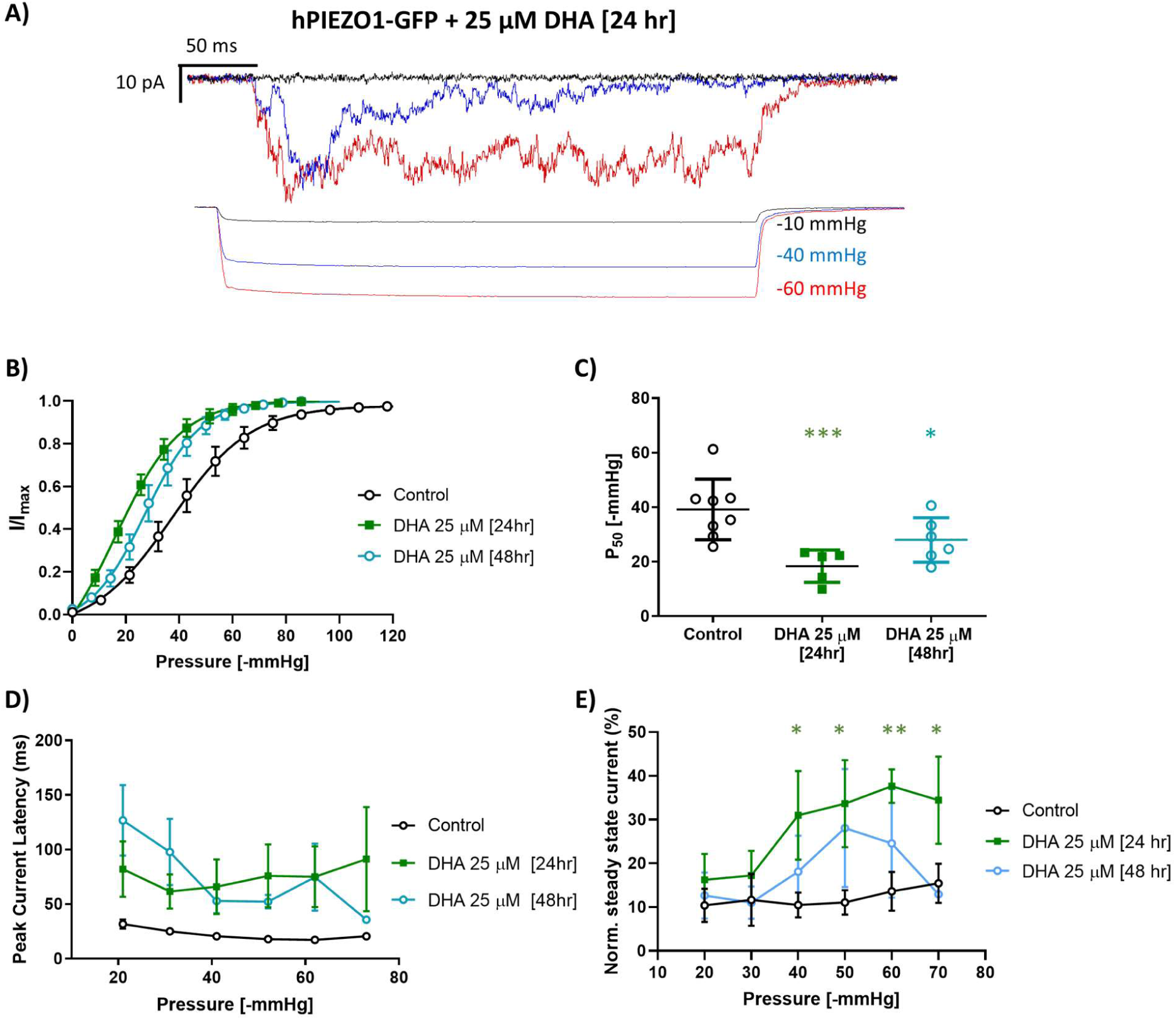
Effect of DHA treatment on PIEZO1-GFP. (A) Patch-clamp recording of PIEZO1-GFP in cell attached mode 24 hr treatment with 25 μM DHA recorded at +65 mV pipette voltage. Each current plot and its respective pressure trace are color-coded. (B) Boltzmann fit of Normalized Current versus Pressure plot of PIEZO1-GFP after 24 hr and 48 hr treatment with 25 μM DHA (Control n = 8 cells. DHA 24hr n = 5 cells, DHA 48hr n = 6 cells). C) Quantification of P_50_ values derived from individual Bolzmann functions analysed in (B). (D) Average peak current latency time recorded in the −20 to −70 mmHg pressure regimes for control and DHA-treated cells (Control n = 8 cells. DHA 24hr n = 5 cells, DHA 48hr n = 6 cells). E) Quantification of PIEZO1-GFP current inactivation represented as normalized steady state current at all pressure between −20 and −70 mmHg applied pressure in control, 24 and 48 hr DHA-treated cells (Control PIEZO1-GFP n = 8 cells, DHA 24 hr n = 5 cells, DHA 48 hr = 3 cells). Statistical analysis was performed using One-Way ANOVA, * P < 0.05, ** P < 0.005, *** P < 0.0005; error bars represent S.E.M.

## DISCUSSION

In this study we have examined the effect of cholesterol depletion on PIEZO1’s response to mechanical stimuli in the form of negative pressure applied to cell-attached membrane patches. Our patch-clamp results indicate that the manipulation of membrane cholesterol alters PIEZO1-mediated mechanotransduction. We observed changes to collective (synchronized) inactivation and a latency of activation of PIEZO1 channel clusters, when MBCD or dynasore was added to stably expressing HEK293T cells (Fig. 1). Rapid inactivation of PIEZO1 by membrane stretch is a key feature of this mechanosensitive ion channel [1, 42]. Slowing inactivation in mutant PIEZO1 channels has been linked to several mechanopathologies – the first being hereditary xerocytosis, a familial anemia [5, 43]. The effect of cholesterol depletion on PIEZO1 has similar characteristics observed for these mutations [43]. For example, two mutations introduced to the ion conducting pore of PIEZO1 resulted in a channel devoid of inactivation[43]. However, it differs because in this case a leftward shift in pressure dependency was seen for the double mutation whereas cholesterol removal caused a rightward shift. At the single channel level both the removal of cholesterol and these point mutations introduced a latency of activation of hundreds of milliseconds, something not present in the wild type channel (Fig.1G and SI Fig.1). Currently, the latency of channels could only be explained by fracture kinetics that predicted the presence of clusters [6]. The association of PIEZO1 with cholesterol-rich membrane domains appears to be a crucial determinant of its mechanotransductive properties [22]. Possible evidence of cluster formation was also observed previously using structure illuminated microscopy (SIM) [18] [44].

Using super resolution microscopy we demonstrate that PIEZO1 indeed forms membrane clusters at the nanoscale. PIEZO1-GFP clusters in the cell membrane have a broad range of sizes (Fig.4), from presumably isolated single channels up to tens of thousands nm^2^ assemblies of the protein. The stable cell line probed here displays a range of PIEZO1-GFP expression levels, as well as a non-homogeneous distribution of the clusters on the cellular basal membrane (Fig.4D). We identified cluster sub-populations that are disrupted and are more sensitive to cholesterol modulation. Approximate estimates of the PIEZO1 protein area based on the most detailed available structure [PDB: 6B3R] suggest that a single trimer projects a ~280nm^2^ area when observed perpendicularly to the membrane plane [10]. Estimating channel numbers in the cluster is complicated by the dome-like structure that is adopted by PIEZO1 and the ability to assess how they would pack [11]. Nevertheless, all the employed treatments appear to reduce the median density of PIEZO1-GFP clusters and to significantly enrich a subset of smaller clusters (~1000-1500 nm^2^ in area) (Fig.4B, histogram). Correlative experiments on live cells allowed us to follow the cluster biophysical properties during treatment (SI Fig.6).

Removal of membrane cholesterol via MBCD appeared to generate more PIEZO1-GFP entities in the membrane (Fig.5B) and to cause a faster diffusion of channels in the basal membrane (Fig.3D, 5D), suggesting that cholesterol domains confine PIEZO1-GFP clusters within lipid boundaries, rather than cytoskeletal fences (SI Fig.4). Interestingly, the MBCD treatment increased the co-localization between PIEZO1-GFP and the raft marker CtxB (Fig.5C), suggesting that upon cholesterol disruption PIEZO1-GFP clusters retained interactions with lipid raft components that are enriched in GM1 ganglioside, the target of CtxB. The association of PIEZO1 and lipid rafts has been suggested since depletion of STOML3 affects PIEZO1 activation [23]. These colocalized PIEZO1-GFP/CtxB entities also appeared to diffuse faster than observed in control conditions (Fig.5D), similarly to what was observed in single-color PIEZO1-GFP in situ imaging (Fig 3D). Our most accurate estimates for the diffusion of PIEZO1-GFP clusters were in the order of 0.01 µm^2^/s, within the same order of magnitude as previously reported by the Pathak lab (0.067μm^2^/s) [45].

In control conditions approximately 40% of detected PIEZO1-GFP entities appeared to be co-localized with ordered domains (Fig. 6E, SI Fig.5), while the remaining fraction co-localized with disordered lipid entities (Fig. 6D). The MBCD treatment did not increase the co-localization of PIEZO1-GFP with disordered lipids (Fig.6E), supporting an interaction between ordered, raft-like membrane fractions and PIEZO1-GFP clusters. Dynasore treatment, which disrupts and reorganizes cholesterol rich domains [46], affected response latency and de-synchronized inactivation similarly to MBCD (Fig.1 G-I), without decreasing PIEZO1-GFP mechanosensitivity (Fig.1 F). Dynasore treatment also failed to increase the diffusion of PIEZO1-GFP clusters in the membrane (Fig.5D) and did not change the degree of colocalization with the marker CtxB when compared to control (Fig.5C). This suggested that, after dynasore treatment, the cell membrane retained intact cholesterol-rich domains able to confine PIEZO1-GFP diffusion observed for the control. In some experiments the treatment also caused an increase in detectable PIEZO1-GFP entities suggestive of cluster breakdown (Fig.6A). Interestingly, dynasore increased the number of disordered Laurdan entities (Fig.6C) while promoting the co-localization of PIEZO1-GFP clusters with disordered lipid entities (Fig.6E). In this case, PIEZO1 showed no increased retention of CtxB (Fig.5C) suggesting that an increased fraction of channel clusters became associated with disordered lipid domains while being separated from cholesterol-rich membrane fractions.

Cholesterol enrichment did not appear to increase the fraction of PIEZO1-GFP in proximity of ordered lipids (Fig.6D) or CtxB (Fig.5C), but significantly increased the fraction of ordered lipids (Fig.6B), indicating that cholesterol enrichment induced an enlargement of existing cholesterol domains. Also, cholesterol enrichment didn’t sensitize the PIEZO1-GFP response or affect its kinetics (Fig.1B), indicating the additional membrane stiffening due to ordered lipids did not affect the channel response. In contrast, PIEZO1 has been reported to be desensitized following membrane stiffening by supplementation of saturated fatty acids [20] which become incorporated into membrane phospholipids. It appears therefore that PIEZO1 sensitivity is mostly affected by the stiffness of non-raft membranes.

To confirm this, we tested whether the membrane fluidity of the disordered fraction of the lipid membrane sensitizes the mechanosensitive response of PIEZO1-GFP. When cultured with DHA, PIEZO1-GFP expressing cells displayed lower activation thresholds (Fig.7B, 7C) and an increased latency of the response (Fig.7D, SI Fig.1). In contrast with a recent study employing whole-cell patch-clamping and activation of PIEZO1 by cell indentation [20], our cell-attached patch data revealed a significant sensitization to applied suction after 24 and 48 hours of PUFA enrichment (Table 1). The activation thresholds quantified in Table 1 are in good agreement with previously published estimates [18] [43].

This suggested that the higher line tension [47, 48] of the disordered membrane phase lowered the threshold for channel activation (Fig.1F), while a more fluid lipid medium delayed the propagation of a tension wave across the patched membrane, as seen in MBCD and dynasore experiments (Fig.1G, SI Fig.1). Interestingly, DHA-treated cells displayed a dose-dependent PIEZO1-GFP inactivation response causing a reversible loss of inactivation at high pressure regimes (Fig.7E), previously reported to occur independently of the magnitude of the stimulus [20]. This is consistent with previous observations that loss of inactivation in PIEZO1 can also be induced by repeated mechanical stimulation causing synchronous loss of inactivation of all channels in a membrane patch [43]. The previous observations that inactivation is lost after repeated or excessive stretch might be due to kinetic de-mixing of cholesterol rich domains in the membrane patch which is compatible with the idea that the channels exist in clusters [49-51].

We envision a scenario based on the “picket-fence model” [52] where cholesterol fences, rather than cytoskeletal fences, organized as a dynamic network on the cell surface, delimit the 2D diffusion of PIEZO1 clusters. A diagram depicting a possible mechanism of direct cholesterol action on mechanosensitive properties of PIEZO1 is shown in Fig. 8. In our view stiffened lipid structures connecting several clusters would allow a fast propagation of membrane tension between them facilitating the synchronized gating behavior of the channels. Furthermore, here we also show that sensitization of the channel response can be achieved by ‘softening’ the non-raft phase of the membrane (Fig.7B, 7C) rather than by further ‘stiffening’ the raft phase (Fig.1B-F). The canonical PIEZO1 response therefore appears to be dependent on cohesion with cholesterol-rich domains to collectively inactivate, and non-raft domain lipid composition to regulate their sensitivity to tension.

**Figure 8.**
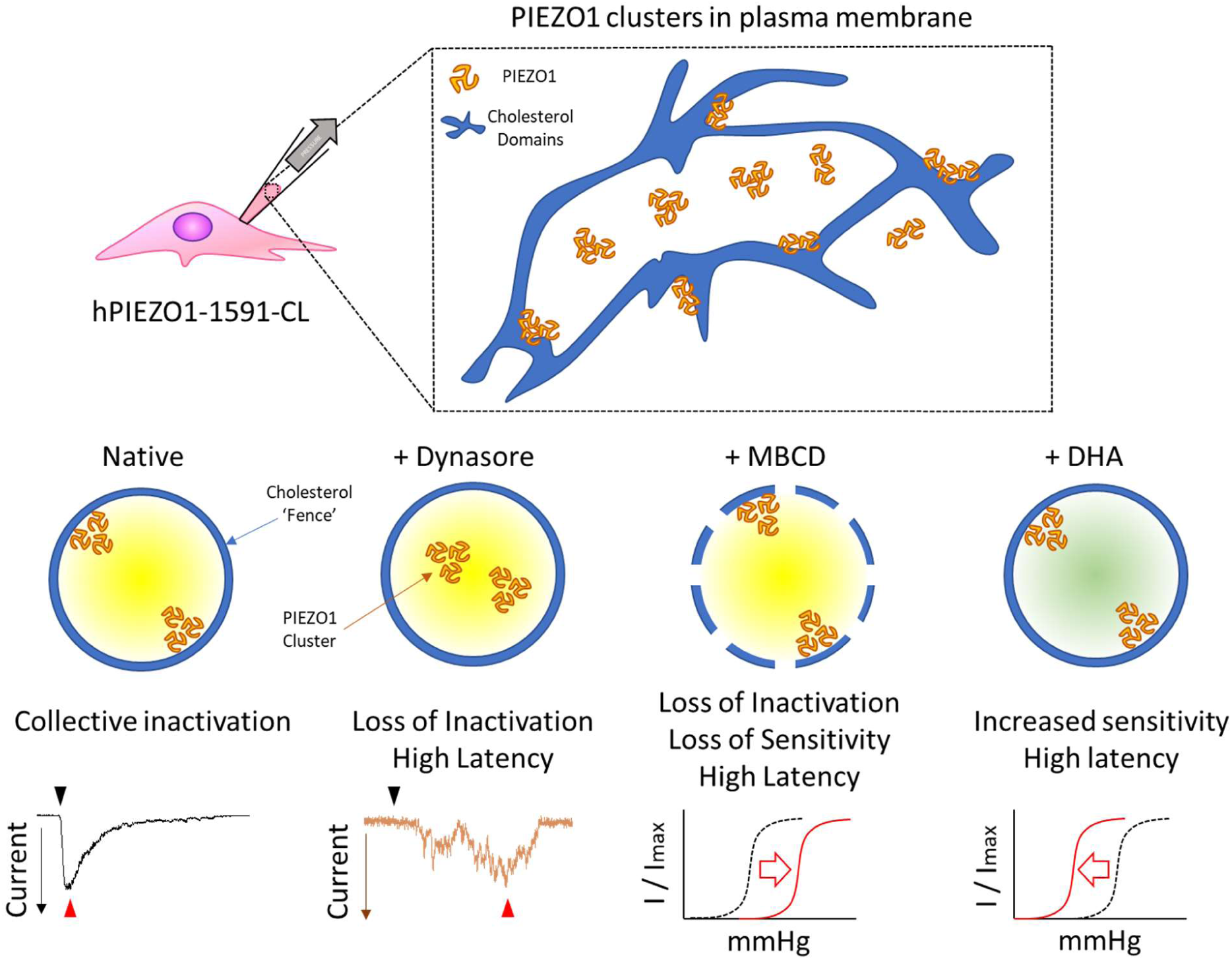
Association between PIEZO1 and cholesterol domain is a major determinant of channel kinetics. Schematic model of the distribution of PIEZO1 clusters in the plasma membrane in relation to cholesterol-rich domains, or ‘fences’. PIEZO1 clusters residing within intact cholesterol-rich domains readily respond to membrane stretch and inactivate in a coordinated fashion. Cholesterol enrichment increases the extent of the cholesterol-fences without compromising the PIEZO1 response. Decoupling of PIEZO1 domains from cholesterol domains results in loss of collective inactivation of channel clusters while loss of structural integrity of such domains results in decreased pressure sensitivity. Membrane-dependent inactivation represents a separate inactivation mechanism which differs from the inherent Piezo1 activation as a property of the channel protein itself [43, 49]. Incorporation of poly-unsaturated fatty acids into membrane phospholipids sensitizes the channel to applied pressure, suggesting that the mechanical properties of the disordered lipid phase is a crucial factor in determining PIEZO1 mechanosensitivity.

Finally, the modulation of cholesterol-dependent inactivation was also observed for the MBCD-treated PIEZO1 R2456H mutant (Fig.2B, 2C). Loss of inactivation after MBCD treatment was observed in both transiently overexpressed WT PIEZO1 (SI Fig.3C) and stably expressing PIEZO1-GFP cells (Fig.1I). However, only the former appeared to be experiencing a concomitant change of sensitivity (Fig.1F). We speculate that transient (24-48hr) overexpression of the channel wasn’t sufficiently long to allow proper localization of the PIEZO1 channel with respect to cholesterol domains. In contrast, a stable cell line would have regulated its PIEZO1 levels and subcellular distribution over many generations to avoid long-term toxicity due to acute overexpression, allowing for a significant MBCD-driven modulation of both inactivation and sensitivity to be detected. This conclusion is supported by MBCD effect on N2A cells (SI Fig.2)

The relationship between PIEZO1 sensitivity and propensity to inactivate is complicated by our lack of knowledge of the full channel gating mechanism. PIEZO1 R2456H displays a lower activation threshold (Fig.2E-F) besides its reduced inactivation (Fig.2C), in contrast with previous observations in whole-cell patch-clamp experiments [20, 53]. In this study we confirm that the lipid bilayer participates in the process of inactivation, as previously shown [20], and can exacerbate the non-inactivating mutant phenotype. This suggets during the channel gating cycle, a lipid-dependent component plays a major role in regulating the kinetics of inactivation. This fits well with recent extensive data from the Vasquez lab [20] and extends the effects to cholesterol. These lipid-mediated effects may also explain why native PIEZO1 shows ameliorated inactivation in some cell types [2, 54]. Furtherwork should aim to probe whether these effects are generated by specific lipid-protein interactions, global effects on the mechanics of the bilayer or both. Cholesterol mediated effects on PIEZO1 may be relevant to cell types frequently exposed to cholesterol in the circulation such as red blood cells, vascular endothelial cells [55], leukocytes [56] or cells displaying high levels of membrane cholesterol such as myelinating oligodendrocytes and Schwann cells [57].

## CONCLUSION

Our electrophysiology results suggest that the inactivation of PIEZO1 currents could be mediated by cholesterol domains bridging PIEZO1 clusters and not only an intrinsic effect of the conformational changes of the PIEZO1 protein during its gating cycle, as shown by previous mutation studies [35-37, 43, 53]. This ability of lipids to modulate or even reverse the slow-inactivating phenotype of PIEZO1 mutants has recently been suggested [20]. Along with this study it highlights the critical importance of studying PIEZO1 and its disease causing variants in native PIEZO1 expressing cells where possible.

## SUPPLEMENTARY INFORMATION

**SI Figure 1.**
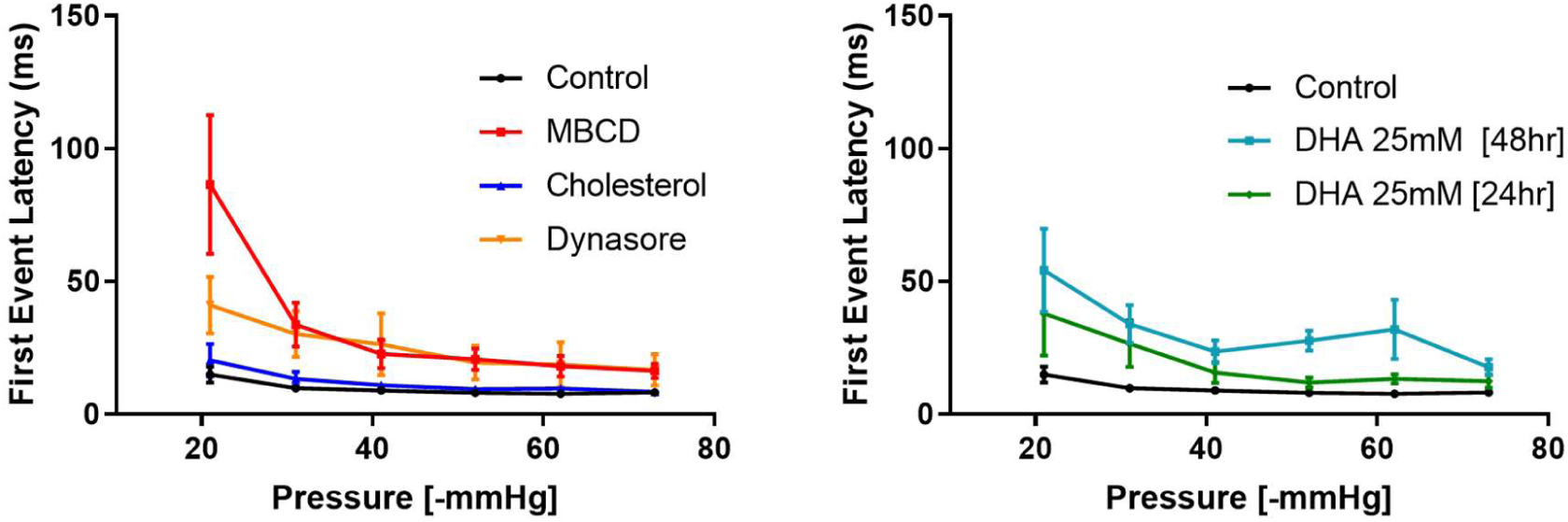
First Event latency of PIEZO1-GFP response. The latency time (ms) was calculated as the time gap from the onset of a computer-controlled High Speed Pressure Clamp negative pressure pulse and the first visible single PIEZO1-GFP response. The PIEZO1-GFP latency was measured during 350 ms-long square suction pulses, ranging from −20 mmHg to −70 mmHg. (Error bars represent S.E.M.)

**SI Fig. 2.**
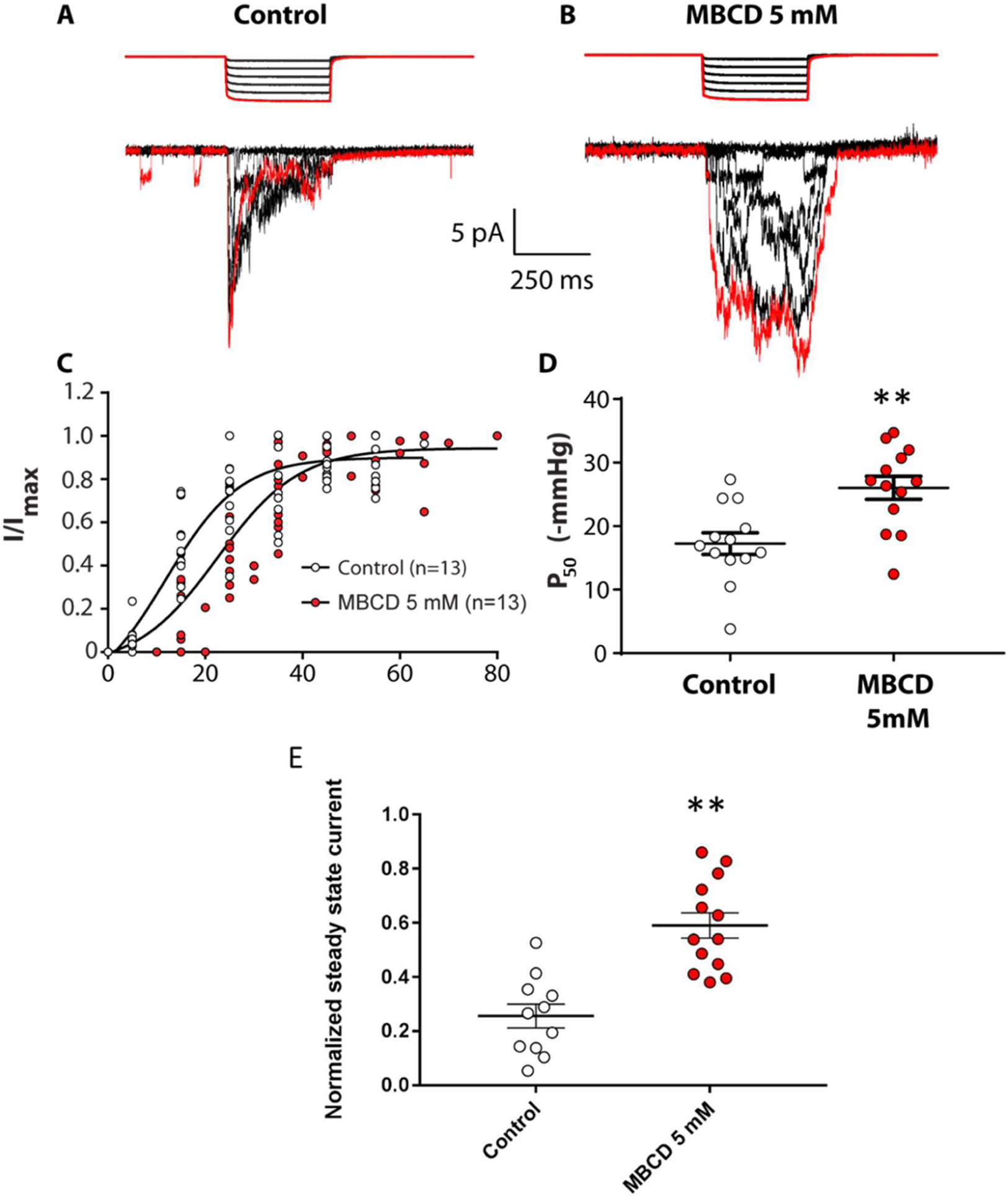
Cholesterol depletion via MBCD in N2A cells. N2A cells were cultured in 10% Fetal Bovine Serum-supplemented Dulbecco’s Modified Eagle’s Medium and treated with 5mM MBCD in serum-free media for 30min at 37’C before experiments. Electrophysiological recordings were collected as described in Methods. A) Computer-controlled High-Speed Pressure Clamp square pressure pulses were applied to untreated N2A cells in cell-attached mode at +65mV pipette potential. Mechanically-evoked currents from native PIEZO1 proteins are shown below the pressure trace. The first pressure pulse and its respective PIEZO1 current trace are highlighted in red. B) MBCD-treated N2A cells display a loss-of-inactivation phenotype when compared to control. C) Combined normalized I/I_max_ values from all control and MBCD-treated cells in the experiment were fitted with a single Boltzmann curve per condition (see Methods). D) Average P_50_ values obtained from Boltzmann fits for each individual cell analyzed under control and MBCD conditions. E) Comparison of normalized steady-state mechanosensitive currents from N2A cells in control and MBCD-treated conditions. (Control n =13, MBCD n= 13). (Student’s T-test, * P < 0.05; Error bars represent S.E.M.).

**SI Fig. 3.**
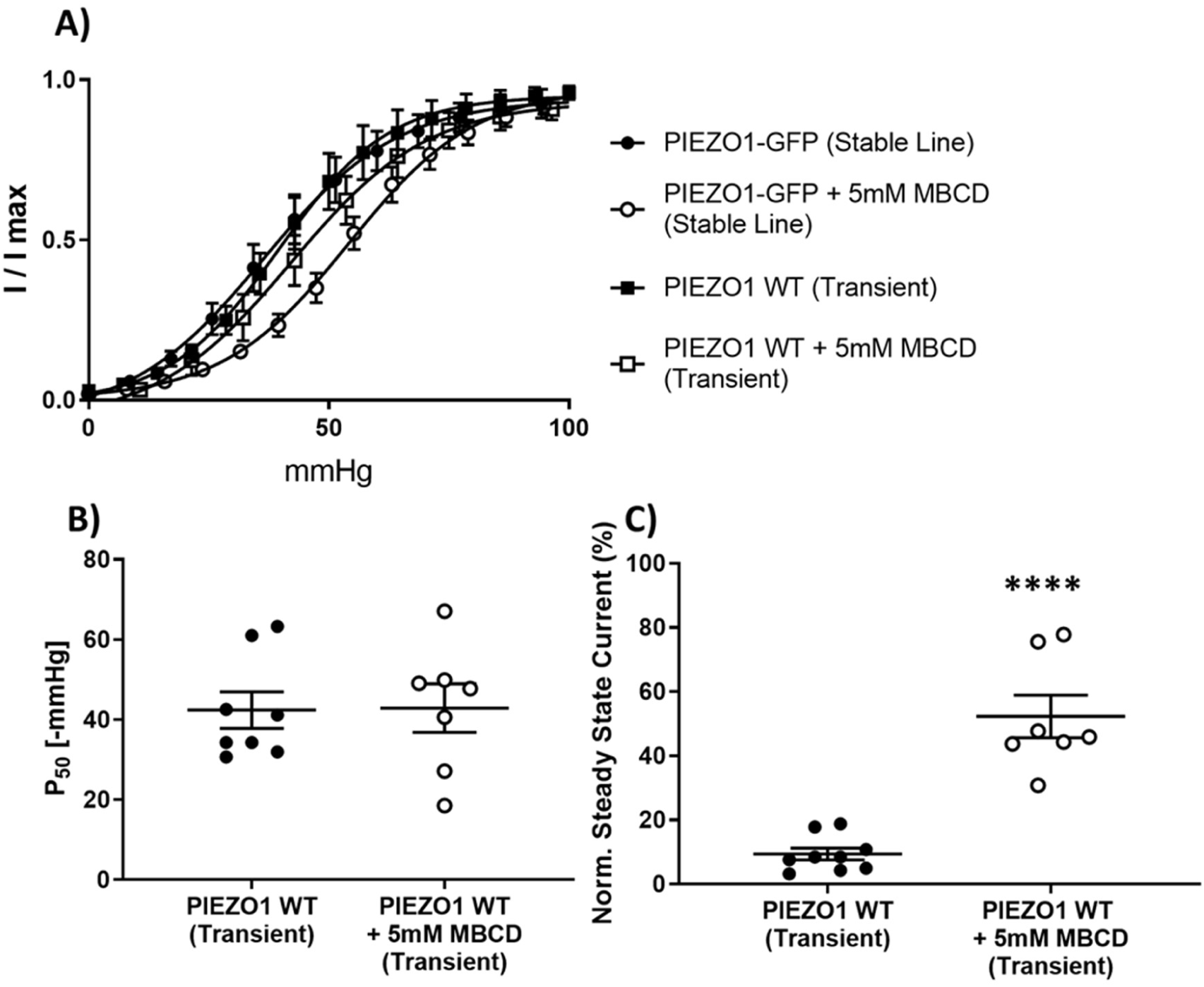
Effect of cholesterol depletion via MBCD on transiently transfected WT human PIEZO1. A) Boltzmann curve fits of normalized I/I_max_ values from all control and MBCD-treated cells stably or transiently expressing human PIEZO1 proteins. Data for the stable cell line is quantified in Fig.1F in the main text. B) Quantification of midpoint activation pressure for transienly expressed WT human PIEZO1 [PIEZO1 WT (n = 8 cells), PIEZO1 WT + MBCD (n = 7 cells)]. C) Quantification of channel inactivation expressed as normalized steady state current for transienly expressed WT human PIEZO1 [PIEZO1 WT (n = 9 cells), PIEZO1 WT + 5 mM MBCD (n = 7 cells)]. Statistical analysis was performed using Student’s T-test (**** P < 0.0001).

**SI Figure 4.**
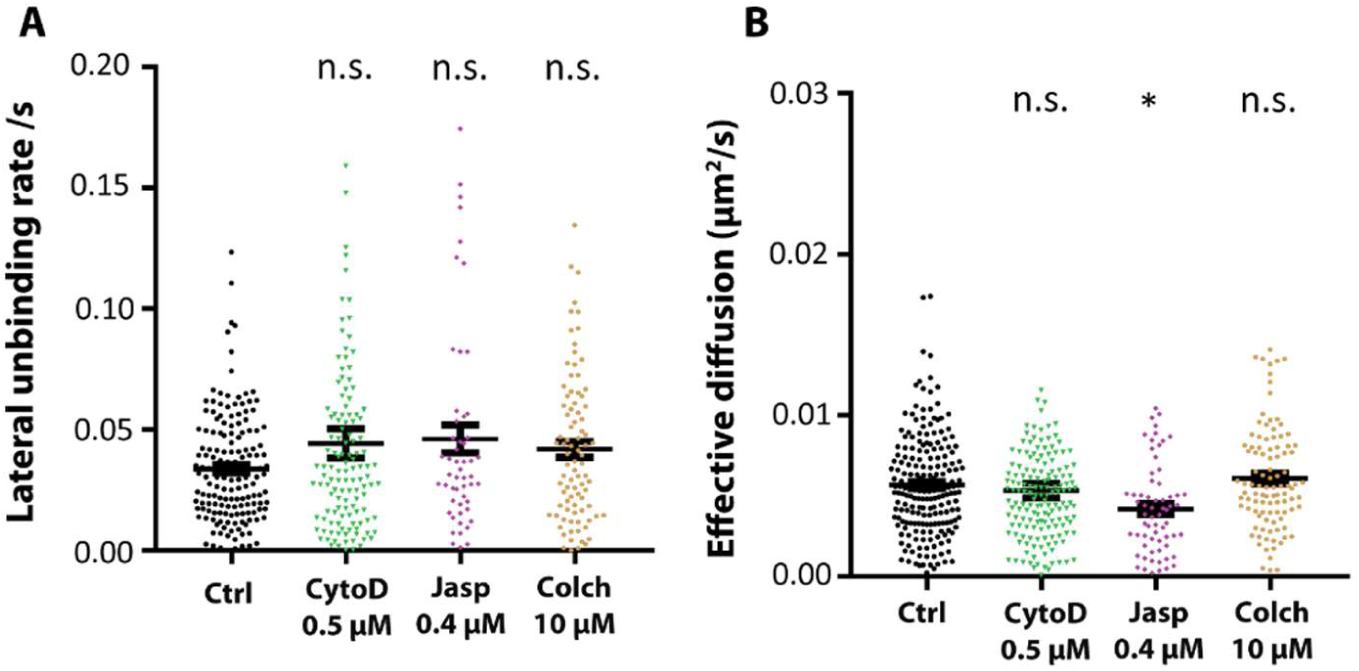
Effect of cytoskeleton manipulation on the membrane diffusion of PIEZO1-GFP. A) Quantification of lateral unbinding rate (s^−1^) of PIEZO1-GFP entities in control and after treatments. B) Effective diuffusion (μm^2^/s) of PIEZO1-GFP entities in control and after treatments. Disruption of the cellular cytoskeleton has no effect on the dispersion rate and the lateral unbinding rate of PIEZO1-GFP clusters while reinforcement reduces its micron-scale diffusion rates. Table legend: Cytochalasin D – CytoD; Jasplakinolide – Jasp; Colchicine - Colch. (One-Way Anova, * P < 0.05; Error bars represent S.E.M.)

**SI Figure 5.**
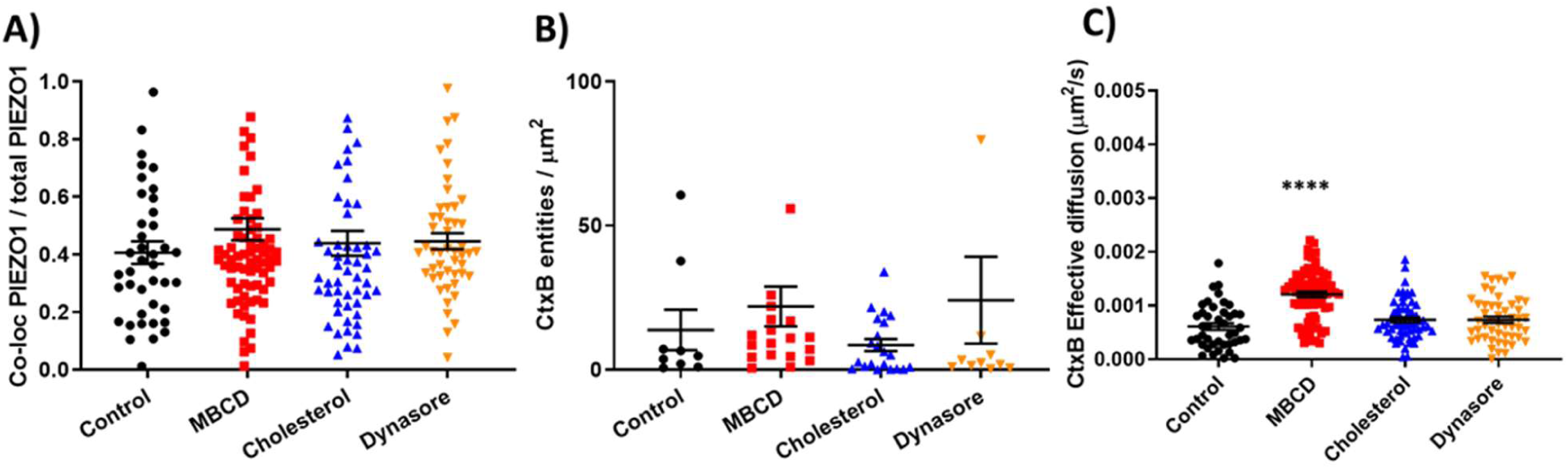
Additional parameters calculated from image Cross Correlation Spectroscopy for assessment of PIEZO1-GFP/CtxB-A555 co-localization in the plasma membrane. (A) Fraction of total PIEZO1-GFP co-localized with CtxB-Alexa555. B) Density of CtxB-Alexa5 entities co-localized with PIEZO1-GFP. C) 2D effective diffusion of CtxB-Alexa555 in the basal plasma membrane. (One-Way ANOVA, **** P < 0.0001; error bars represent S.E.M).

**SI Fig. 6.**
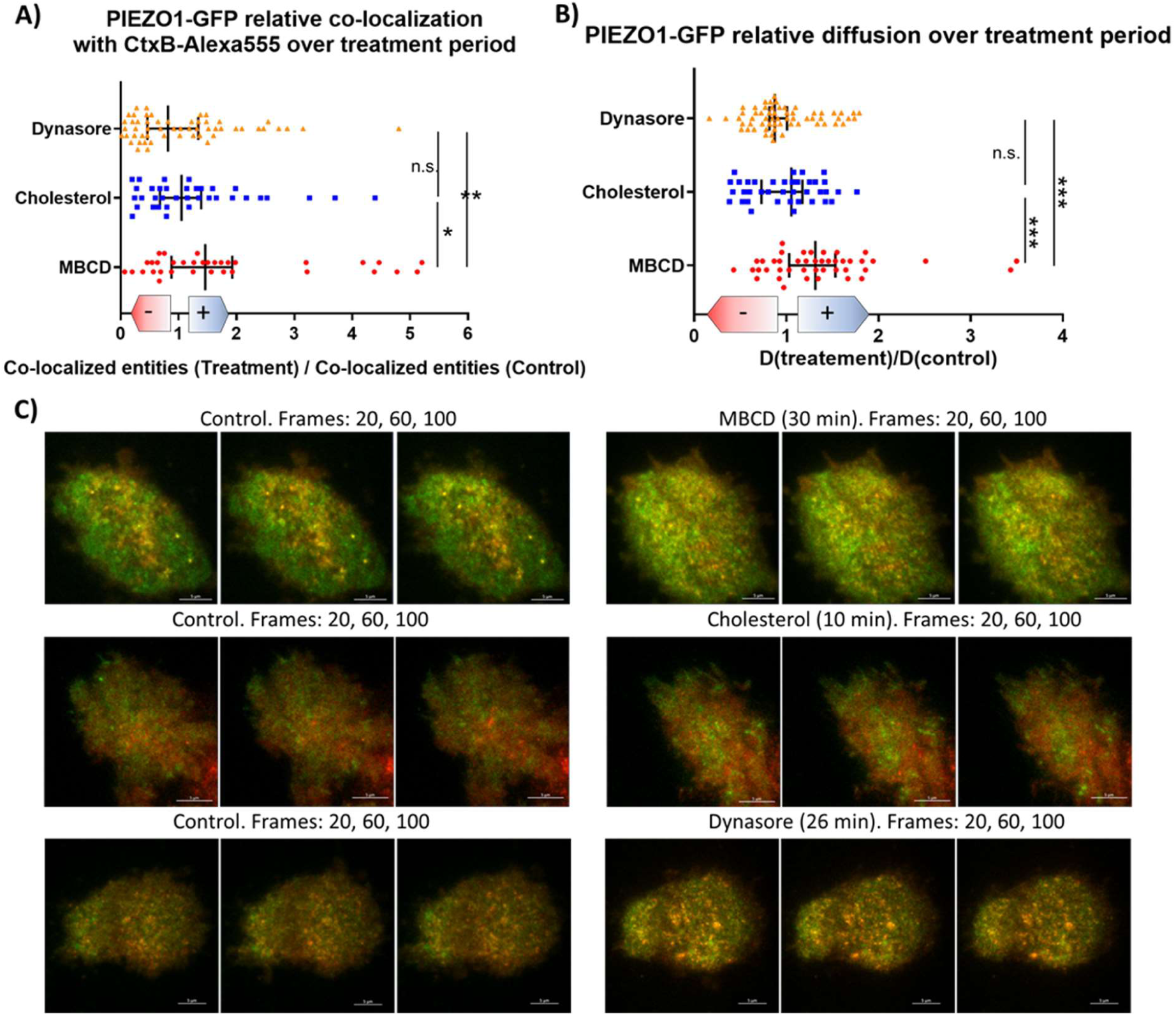
Time lapse imaging of single cells exposed to treatments. Live hP1-1591-GFP cells were stained with CtxB-Alexa555 and imaged periodically in 100-frame time lapses for up to ~70 min. We monitored the amount of co-localized PIEZO1-GFP/CtxB-A555 entities (A) and the diffusion dynamics of PIEZO1-GFP (B) at the single cell level in response to the various treatments used in this study. The results are quantified above and expressed as relative fold-change (“+” shift for increase, “-” shift for decrease) with respect to values recorded prior to drug exposure. Each data point represents a single cell. (C-D-E) Representative frames of image series collected before (left) and after (right) the start of the treatment (from top to bottom: 5 mM MBCD, 50 µg/ml Cholesterol, 80 µM Dynasore). Statistical assessment was performed using One-way anova with multiple comparisons. (* P < 0.05; ** P < 0.005; *** P < 0.0005; Bars indicate Median ± 95% Confidence Interval).

**SI Fig. 7.**
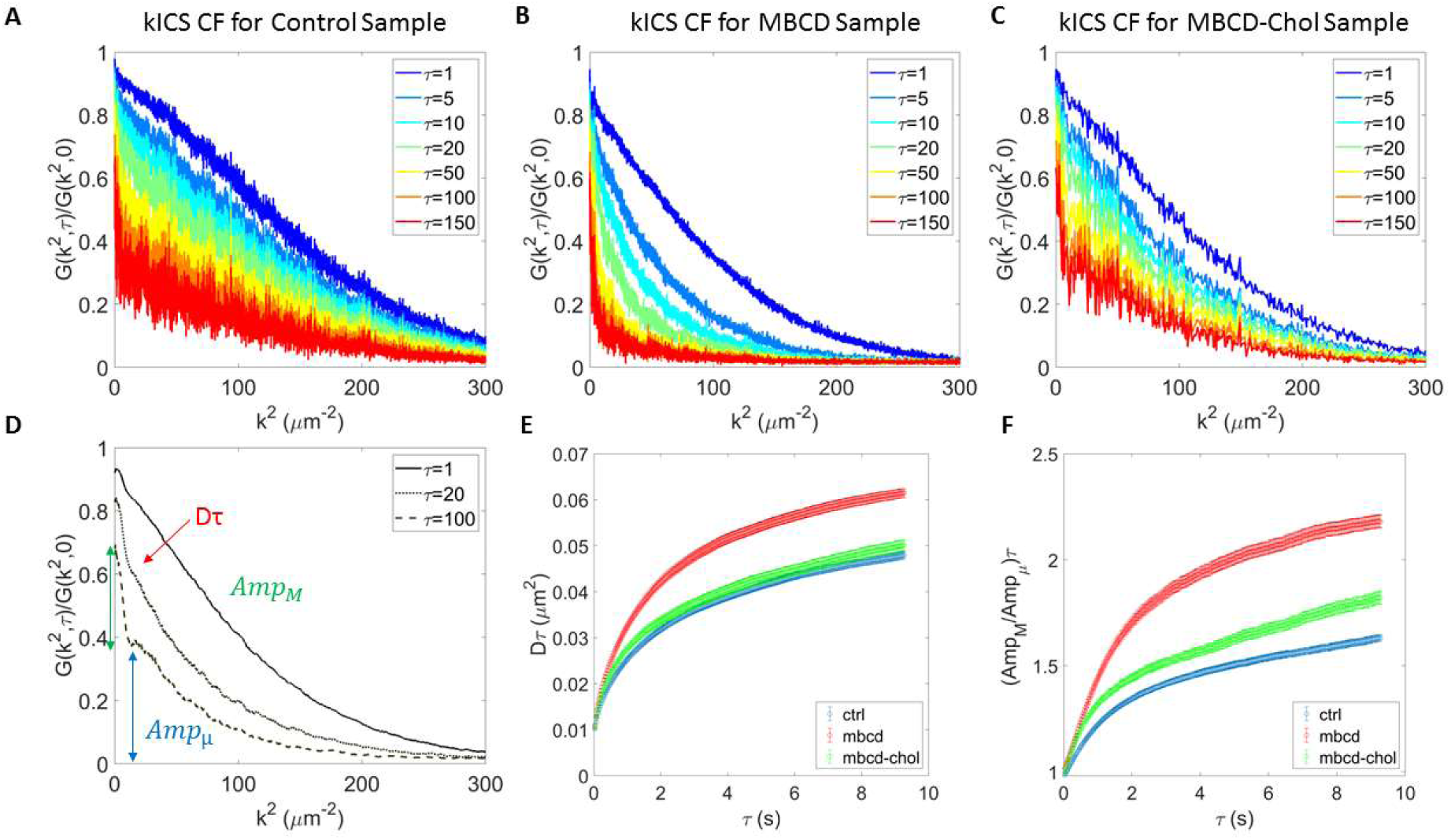
Examples of kICS Correlation Functions. kICS Correlation Functions for Control (A), MBCD (B) and MBCD-Cholesterol (C) conditions, showing effective two dynamic populations more present in Control and MBCD-Cholesterol conditions, while MBCD seemed to remove large k^2^ or in domain diffusive component. (D) From the early decay of kICS CF, at each temporal lag τ, we extracted the effective diffusion decay Dτ, while the amplitude of the two components, macro and micro, Amp_M_ and Amp_μ_ respectively, were extracted from fitting exponential decay vs k^2^, at two different k^2^ ranges. (E) From slope of Dτ vs τ we obtained effective diffusion coefficient, while from late slope of Amp_M_/Amp_μ_ vs τ (F), we obtained the lateral unbinding rate from cholesterol domains. Data in E and F are averages from ~30 cells (>3 experiments) and SEM for 3 conditions teste: Control (blue symbols), MBCD (red) and MBCD-Chol (green).

